# The Crystal Structures of Bacillithiol Disulfide Reductase YpdA Reveal Structural and Functional Insight into a New Type of FAD-Containing NADPH-Dependent Oxidoreductases

**DOI:** 10.1101/2020.12.02.408641

**Authors:** Marta Hammerstad, Ingvild Gudim, Hans-Petter Hersleth

## Abstract

Low G+C Gram-positive Firmicutes, such as the clinically important pathogens *Staphylococcus aureus* and *Bacillus cereus,* use the low-molecular weight (LMW) thiol bacillithiol (BSH) as a defense mechanism to buffer the intracellular redox environment and counteract oxidative stress encountered by human neutrophils during infections. The protein YpdA has recently been shown to function as an essential NADPH-dependent reductase of oxidized bacillithiol disulfide (BSSB) resulting from stress responses and is crucial in maintaining the reduced pool of BSH and cellular redox balance. In this work, we present the first crystallographic structures of YpdAs, namely from *S. aureus* and *B. cereus.* Our analyses reveal a uniquely organized biological tetramer; however, the monomeric subunit has high structural similarity to other flavin disulfide reductases. The absence of a redox active cysteine in the vicinity of the FAD isoalloxazine ring implies a new direct disulfide reduction mechanism, which is backed by the presence of a potentially gated channel, serving as a putative binding site for BSSB in proximity to the FAD cofactor. We also report enzymatic activity for both YpdAs, which along with the structures presented in this work provide important structural and functional insight into a new class of FAD-containing NADPH-dependent oxidoreductases, related to the emerging fight against pathogenic bacteria.

---

Low-molecular weight (LMW) thiols are involved in many important cellular processes in all organisms, including a critical protective role in cells where they maintain cytosolic proteins in their reduced state. They also function as thiol cofactors of many enzymes in scavenging of e.g. reactive oxygen species (ROS), reactive chlorine species (RCS), and reactive electrophilic species (RES), and in detoxification of toxins and antibiotics. LMW thiols are also involved in protection against heavy metals and in metal storage^1–2^. The LMW thiol glutathione (GSH; γ-glutamyl-cysteinylglycine), produced in most eukaryotes, Gram-negative bacteria, and some Gram-positive bacteria, is the most abundant LMW thiol antioxidant, contributing to the control of redox homoeostasis. In redox reactions, GSH is continuously oxidized to glutathione disulfide (GSSG), which can be rapidly converted back to GSH by glutathione reductase (GR)^3^, to maintain the required GSH/GSSG ratio that is important for the cellular redox balance. Furthermore, an important post-translational modification for regulating protein function and protecting exposed cysteine residues from irreversible oxidative damage is the reversible formation of GS-S-protein disulfides (S-glutathionylation) on proteins^3^. Although GSH is the predominant LMW thiol in eukaryotes and Gram-negative bacteria, most Gram-positive bacteria utilize other, distinctly different LMW thiols. High G+C content Gram-positive bacteria (Actinobacteria) produce mycothiol (MSH; AcCys-GlcN-Ins)^4^, whereas low G+C Gram-positive bacteria (Firmicutes) produce bacillithiol (BSH; Cys-GlcN-Mal)^5–6^ (Scheme 1A), serving analogous functions to GSH. BSH is produced by several clinically important human pathogens including *Staphylococcus aureus* (*Sa*), *Bacillus cereus* (Bc), *Bacillus subtilis* (*Bs*), and *Bacillus anthracis (Ba)^5^.* BSH and derivatives such as N-methyl-bacillithiol (N-Me-BSH) have been suggested to be the most broadly distributed LMW thiols in biology^7^.

During infection, many pathogens encounter human neutrophils and macrophages capable of generating ROS and RCS. Also, they are frequently exposed to RES as secondary oxidation products from ROS and RCS as well as from external sources, such as antibiotics^8^. To limit the extent of damage, Firmicutes, such as *Sa*, rely on mechanisms involving BSH as important strategies to combat these toxic and reactive species during infection^9–13^. Another important role of BSH is protein thiol protection through S-bacillithiolation, analagous to S-glutathionylation in eukaryotes^8, 13–18^. De-bacillithiolation of proteins is catalyzed by bacilliredoxins (Brxs).

Analogous to glutaredoxins (Grxs), Brxs attack the active site Cys on the BSH-mixed protein disulfide on S-bacillithiolated substrates, transfering BSH to the Brx active site Cys. The Brx-SSB intermediate is reduced by BSH, leading to oxidized bacillithiol disulfide (BSSB). Alternatively, BSH can react directly with ROS, again leading to oxidation of BSH to BSSB^19^ (Scheme 1). While GSSG is recycled by GR, a recent study showed that the flavoenzyme YpdA from *Sa* consumes NADPH^20^, and another confirmed that *Sa* YpdA reduces BSSB under aerobic conditions^21^. Evidence that BSSB is recycled by the FAD-containing NADPH-dependent disulfide oxidoreductase YpdA, which along with BrxA/B and BSH biosynthesis enzymes BshA/B/C is only present in BSH-containing bacteria, has provided insight into the understanding of the Brx/BSH/YpdA pathway and the recycling of BSSB to maintain the reduced BSH pool (Scheme 1B). However, many questions have remained unanswered regarding YpdA and BSSB reduction. Are YpdA orthologs from other Firmicutes able to reduce BSSB? How structurally similar are YpdAs to other flavoenzymes? How is the mechanism for BSSB reduction in YpdAs? These questions are the focus of this investigation.

In this work, we aimed to investigate whether YpdA acts as a common BSSB reductase by examining a putative YpdA from another Firmicute, *Bc*. *Sa* and *Bc* YpdAs were expressed and purified to measure their enzymatic activity through the consumption of NADPH (340 nm). The enzymes showed similar consumption rates for reactions with and without BSSB substrate present, indicating that the enzymes are highly oxygen sensitive and are reoxidized in the presence of oxygen (Figure S1). This oxygen sensitivity is consistent with previous work on related ferredoxin/flavodoxin NAD(P)^+^ oxidoreductases (FNRs)^22–23^. Therefore, to specifically measure BSSB reduction, enzymatic assays were performed under strict anaerobic conditions (Figure 1). This confirmed that both *Sa* and *Bc* YpdAs are able to reduce BSSB with an NADPH consumption in the same order of magnitude as substrate added to the reactions. With respect to other possible functions of YpdA, previous work demonstrated that *Bc* YpdA has only a limited activity towards flavodoxins (Flds), and the Fld-like protein NrdI^22–23^, strengthening the notion of YpdAs role as a BSSB reductase, and not an FNR, in Firmicutes.

**Figure 1:**
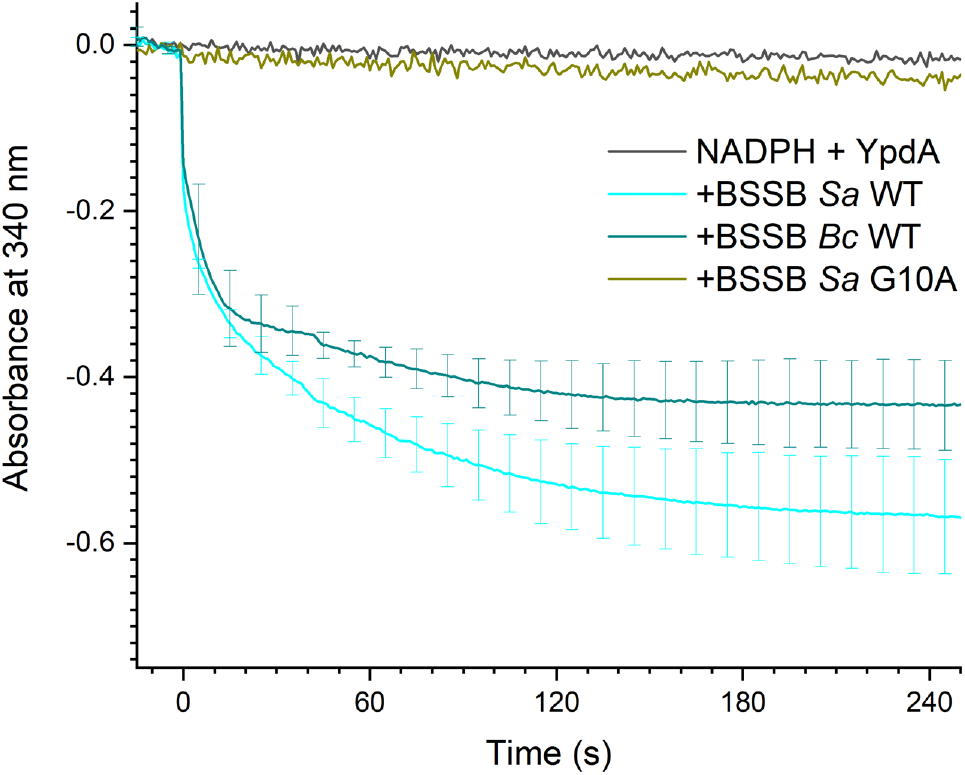
Enzymatic activity of YpdAs as BSSB reductases. YpdA from *Bc* and *Sa* are both able to reduce BSSB, as seen from the NADPH consumption under anaerobic conditions. The *Sa* YpdA G10A mutant has no enzymatic activity towards BSSB.

Although important functional discoveries in the understanding of YpdA as a flavin disulfide reductase have been made, structural information and details offering insight into the YpdA reaction mechanism have been missing. Here, we present the two first reported crystal structures of YpdA; the homologous *Bc* YpdA (1.6 Å resolution) and *Sa* YpdA (2.9 Å resolution) (Figure 2 and Table S1), providing an important missing link in the understanding of a vital redox pathway in Firmicutes.

**Figure 2:**
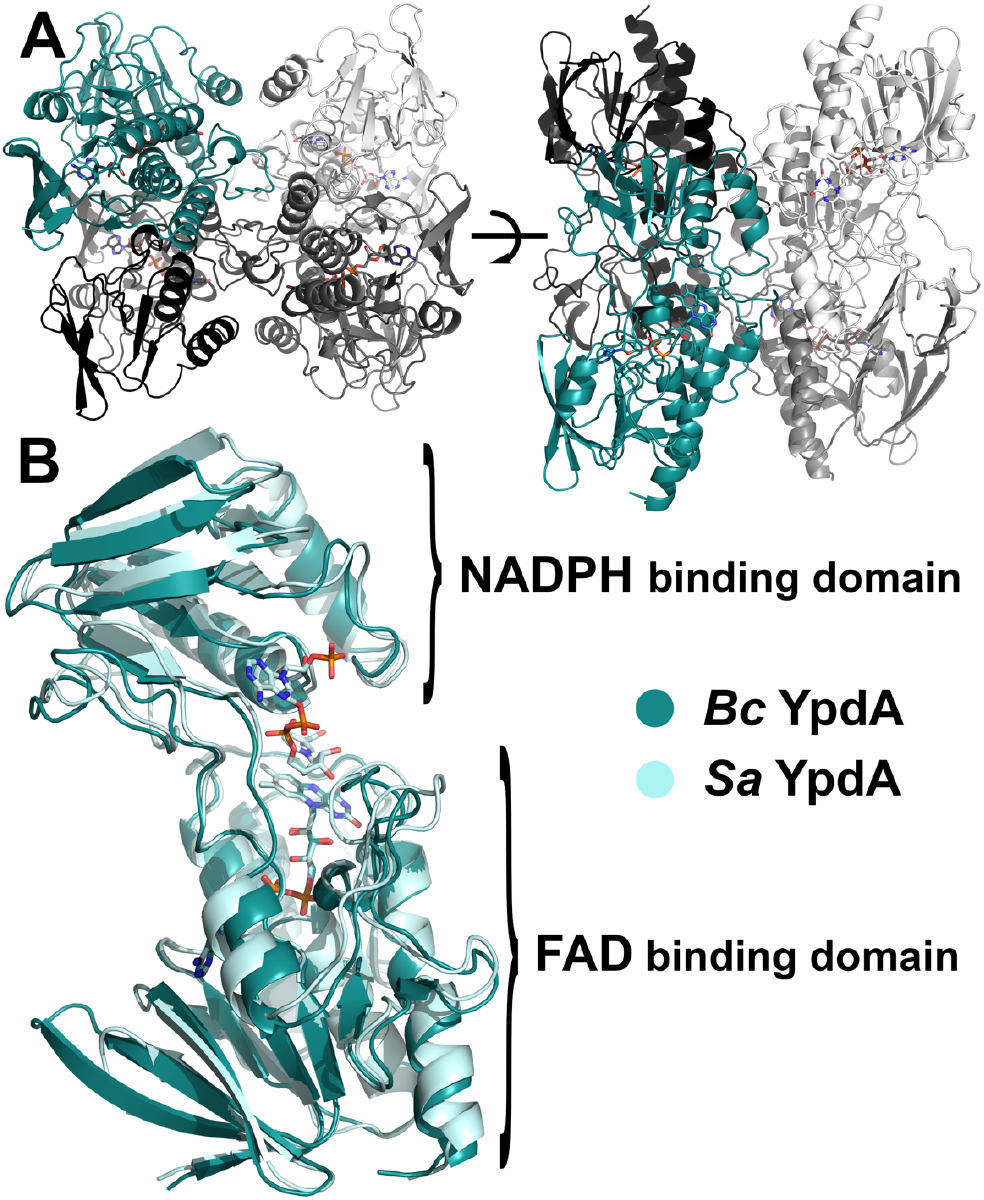
Crystal structures of *Bc* and *Sa* YpdA. (A) Overall structure of the *Bc* YpdA tetramer, seen from two different orientations, colored by chain. (B) Monomer structure alignment of *Bc* and *Sa* YpdA, displaying the NADPH and FAD binding domains. Cofactors are represented as sticks and colored by atom type.

The monomeric subunits of YpdA (Figure 2) show, as expected, high structural similarity to members of the “two dinucleotide binding domains” flavoproteins (tDBDF) superfamily with NADPH and FAD binding domains containing the three-layer ββα sandwich Rossman-like folds^24^. The structural similarity was confirmed by a DALI search with RMSD-values in the range of 2.4-4.6 (Table S2), as also shown in the structural and sequence alignments in Figures S2 and S3 (see Sections S3 and S4 in SI). The oligomeric state of YpdA is, however, unique, as both structures comprise a conserved tetrameric core (Figures 2 and S2). The biological tetrameric oligomerization state of YpdA was confirmed through crystal packing (Figure S4), dynamic light scattering (DLS), and native polyacrylamide gel electrophoresis (PAGE) analyses (Figure S5). The overall YpdA tetramer reveals dimer interfaces that are different than those observed in structures of e.g. flavoprotein monooxygenase (FPMO) (PDBid 4C5O) and TrxR (PDBid 5VT3,1TDF) (Figure S2), presenting a unique biological assembly of a new type of FAD-containing NADPH-dependent oxidoreductases. This is further substantiated by phylogenetic analyses on selected Firmicutes, revealing that YpdAs comprise a separate clade and is different to other structurally similar oxidoreductases (Figure S6).

The Rossman-like FAD binding domain of YpdA contains the canonical glycine-rich signature sequence motif GXGXXG/A (G_10_GGPC14G in *Sa* and *Bc*) (Figure S7)^24–25^. A previous study showed that the *Sa* YpdA G10A mutant, a mutation known to disrupt cofactor binding, is unable to consume NADPH under aerobic conditions^20^. Here, we confirm that the *Sa* YpdA G10A mutation results in loss of the FAD cofactor, and hence, rendering YpdA in its inactive apo-form (Figure S8), unable to consume NADPH under aerobic or anaerobic conditions, and incapable of reducing BSSB (Figure 1). Furthermore, it was recently suggested that YpdA acts on BSSB through a conserved residue (Cys14). This was based on the ceased enzymatic activity in a YpdA C14A mutant, implying that Cys14 acts as an active site residue in YpdA for BSSB reduction^21^. The YpdA crystal structures presented in this work reveal, however, that Cys14 is located in a buried environment, ~8 Å away from the FAD isoalloxazine ring, making reactions with both FAD and BSSB unlikely (Figures 3A and S7). This is in contrast to GR (PDBid 1GRB), where the active site solvent/substrate-accessible cysteine is located 3.4 Å away from the isoalloxazine ring. Generation of potential solvent channels around Cys14 with HOLLOW^26^ shows inadequate space for BSSB entry or binding in the proximity of Cys14, as well as between Cys14 and the FAD cofactor. The notion that Cys14 is unlikely to directly participate in the reaction mechanism is further strengthened by a conserved orientation of secondary structure elements and residues closely lining and shielding the GXGXXG/A motif in all 12 YpdA subunits observed in the asymmetric units of the *Bc* and *Sa* YpdA structures (Figures S7 and S4). Although this Cys is conserved in YpdA homologs in Firmicutes, it is replaced by other conserved residues within YpdAs from other phyla suggested to use BSH or N-Me-BSH (Figures S9 and S10). If these putative YpdAs are to function as BSSB disulfide reductases, Cys cannot be essential for a universal reaction mechanism in YpdAs, even if it could be important for e.g. cofactor binding in Firmicutes. The structures shows that Cys14 points into a confined space (Figure S7), possibly indicating that replacement with a less bulky amino acid, such as in the inactive *Sa* YpdA C14A mutant^21^, could destabilize important protein-cofactor interactions. Future studies need to be performed to investigate putative YpdA homologs from other phyla as potential BSSB reductases to confirm this.

**Figure 3:**
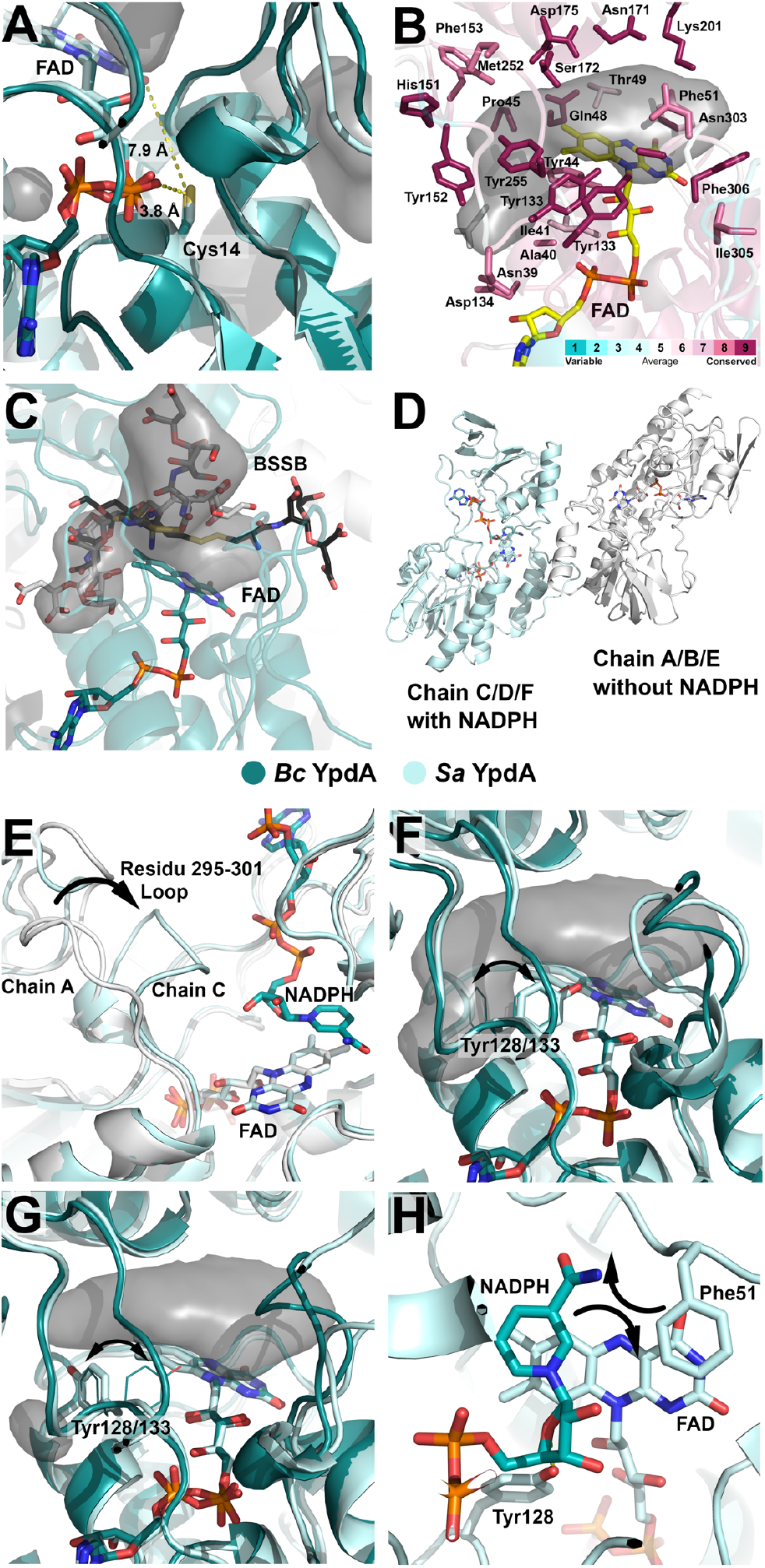
Structural features of YpdA. (A) Cys14 is located in a buried pocket, 7.9Å (*Bc*) away from the FAD isoalloxazine ring. (B) Solvent channel lined by conserved residues, generated with HOLLOW and ConSurf. (C) Potential BSSB binding sites in the HOLLOW-generated solvent channels in proximity to the FAD cofactor. (D) NADPH-bound and NADPH-free states (*Sa* YpdA). (E) Flexible loop involved in entry/binding of NADPH and possibly BSSB (*Sa* YpdA). (F) Potential gating mechanism for BSSB entry by Tyr128 in open conformation and (G) closed conformation (*Sa* YpdA), and (H) different stacking conformations of the NADPH nicotinamide and Phe51 in *Sa* YpdA.

If not through Cys, then where and how does BSSB interact with YpdA? Our crystal structures show a large solvent channel lined by conserved residues spanning the entire monomer on the *re*-face of the isoalloxazine ring of FAD, providing a sufficiently large surface space for BSSB binding (Figures 3B, S11, and S12). Figure 3C presents three potential binding orientations of BSSB fitted within the channel in close proximity to FAD. One is positioned with the BSSB disulfide bond close to the reactive C4a (C4x, FAD numbering) atom of FAD (Figure 3C), and another is protruding slightly into the NADPH binding channel. This clearly demonstrates that BSSB could bind close to FAD for a potential reaction, although there is an overlap between the NADPH and BSSB binding sites. Surprisingly, despite not adding NADPH/NADP^+^ to the crystallization or cryo solutions, we observed electron density for bound NADPH in two of the four subunits of the core *Sa* YpdA tetramer, as well as one out of two additional chains seen in the crystal packing, resulting in dimers of NADPH-bound and NADPH-free states (also observed in an additional *Sa* YpdA crystal structure, unpublished data) (Figures 2, 3D and S13). This could point towards cooperativity and asymmetric enzyme activity in YpdA. NADPH binding is clearly gated by a loop movement (residues 295-301), closing parts of the NADPH binding channel in chains C/D/F as compared to the NADPH-free state (chains A/B/C) (Figure 3E). Further, in the NADPH-bound state (chains C/D/F), a Tyr residue (Tyr128, *Sa* numbering) is hydrogen-bound to the NADPH ribose moiety (open conformation) (Figure S13D), allowing for access of BSSB to the suggested substrate binding channel (Figures 3F). In NADPH-free subunits of YpdA, Tyr128 adopts an alternative conformation (closed conformation), flipping away from the NADPH cofactor with a rotamer orientation that obstructs the solvent channel for BSSB entry/binding (Figure 3G), suggesting that Tyr128 is involved in a potential gating mechanism. Although no electron density is observed for NADPH in the Bc YpdA structure, here, Tyr133 (*Bc* numbering) adopts both conformations (open/closed) in all four chains (chains A-D), supporting a flexible and possibly regulative role in the gating of substrate entry/binding to the YpdA active site. In *Sa* YpdA, the nicotinamide binds in a close stacking orientation above the re-face of the isoalloxazine ring. Also, Phe51 stacks on the *re*-face (Figure 3H). Two conformations of NADPH are observed in chain D, resulting in different nicotinamide stacking orientations above the isoalloxazine ring combined with two different orientations of Phe51. This indicates that Phe51 can flip away from its normal stacking orientation, and potentially, together with Tyr128, function in the gating of substrate entry and binding on the FAD *re*-face.

Our crystal structures have revealed that YpdA lacks an accessible active site cysteine, making a cysteine-based thiol mechanism unlikely. We observe a probable BSSB binding site directly above the isoalloxazine ring, possibly regulated through an amino acid gating mechanism of the channel on the *re*-face of the isoalloxazine ring. Based on our findings, we propose a new, simpler and more direct reaction mechanism for YpdA, resembling the classical flavin disulfide reductase mechanism^27–28^, but without the cysteine-based dithiol step (Scheme 2). First, NADPH (substrate 1) binds to YpdA (*E*^0^’ox/hq = −242 mV, *Bc* YpdA^23^) with the loop 295-301 closing in, Tyr128/133 flips from closed to open conformation, and Phe51 possibly opens up the re-face. Next, in the reductive half-reaction (Δ*E*^0′^ = 82 mV, *Bc*), the FAD group is reduced by NADPH through hydride transfer. NADP^+^ leaves, and BSSB (*E*^0′^_BSSB/BSH_ = −221 mV^29^) (substrate 2) binds close to the reduced isoalloxazine ring likely with the disulfide close to the reactive C4a. Finally, in the oxidative half-reaction (Δ*E*^0′^ = 21 mV, *Bc*), BSSB is reduced to a thiol-thiolate pair where the thiolate near C4a forms a C4a-cysteine adduct with the flavin, ultimately leading to the reduced BSH products. This putative mechanism is consistent with our structural investigations and activity studies; however, further studies giving insight into the details of the mode of action in YpdAs are interesting topics for future investigations.

This study provides the first crystal structures of YpdAs from two homologous species, demonstrating that YpdAs comprise a new class of oxidoreductases and providing structural insight into a new reaction mechanism. Our findings present an important missing link in the field of redox biology and the regeneration of BSSB; a critical process in many clinically important human pathogens. Structural insight into YpdA may provide a new potential target for antimicrobial drug design and the fight against pathogenic bacteria.

## Supporting Information

Experimental procedures used in this study, including protein expression, purification, and characterization; activity measurements; protein crystallography; and bioinformatics and structural analysis. Supplemental Figures S1-S13 and Tables S1-S2.

## Acknowledgement

We thank Dr. B. Dalhus for access to crystallization screening at the Regional Core Facility for Structural Biology at the South-Eastern Norway Regional Health Authority (2015095). We gratefully acknowledge MAX IV and the ESRF for providing beam time and the staff of beamlines BioMax (MAX IV) and ID23-1 (ESRF) for excellent technical assistance. We thank Dr. Tor Erik Kristensen for valuable discussions on the YpdA reaction mechanism. This project has been funded by grants from the Research Council of Norway (231669 and 301584). The research leading to these result has been supported by the project CALIPSOplus under the Grant Agreement 730872 from the EU Framework Programme for Research and Innovation HORIZON 2020. X-ray coordinates and structure factors have been deposited in the PDB Database PDBids: 7A7B (*Sa* YpdA), 7A76 (*Bc* YpdA)).

## Abbreviations

LMW: low-molecular weight;
ROS: reactive oxygen species;
RES: reactive electrophilic species;
RCS: reactive chlorine species;
GSH: glutathione;
GSSG: glutathione disulfide;
MSH: mycothiol;
BSH: bacillithiol;
BSSB: bacillithiol disulfide;
N-Me-BSH: N-methyl-bacillithiol;
*Bc*: *Bacillus cereus*;
*Bs*: *Bacillus subtilis*;
*Ba*: *Bacillus anthracis*;
*Sa*: *Staphylococcus aureus*;
GR: glutathione reductase;
TrxR: thioredoxin reductase;
FPMO: flavoprotein monooxygenase;
FNR: ferredoxin/flavodoxin NAD(P)^+^ oxidoreductase;
Fld: flavodoxin;
Grx: glutaredoxin;
Brx: bacilliredoxin;
BshA/B/C: BSH biosynthesis enzymes A/B/C;
YpdA: bacillithiol disulfide reductase;
tDBDF: two dinucleotide binding domains flavoprotein superfamily;
FAD: flavin adenine dinucleotide;
NADPH: nicotinamide adenine dinucleotide phosphate, reduced form;
NADP^+^: nicotinamide adenine dinucleotide phosphate, oxidized form;
PAGE: polyacrylamide gel electrophoresis;
DLS: dynamic light scattering;
*E*^0′^: standard reduction potential at pH 7.

## Schemes, figures and legends

**Scheme 1:**
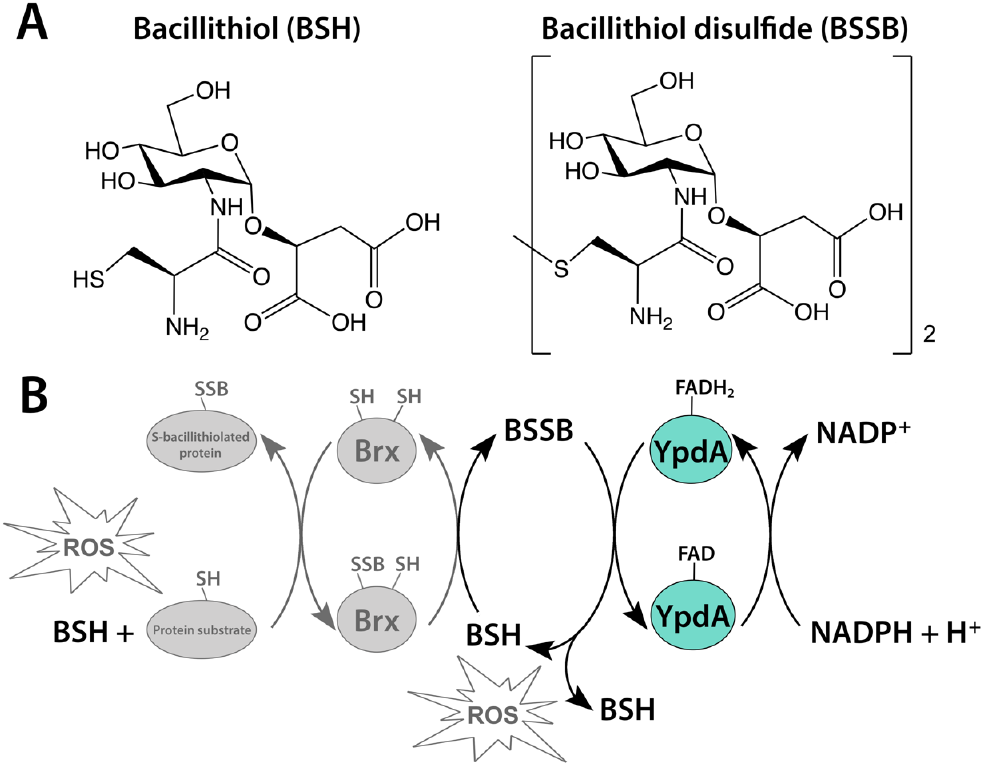
Bacillithiol and the regeneration of reduced bacillithiol by YpdA. (A) Structure of bacillithiol (BSH, left) and bacillithiol disulfide (BSSB, right). (B) Reduction of BSSB by YpdA in the regeneration of BSSB to BSH following protein de-bacillithiolation or ROS-detoxification by BSH in Firmicutes.

**Scheme 2:**
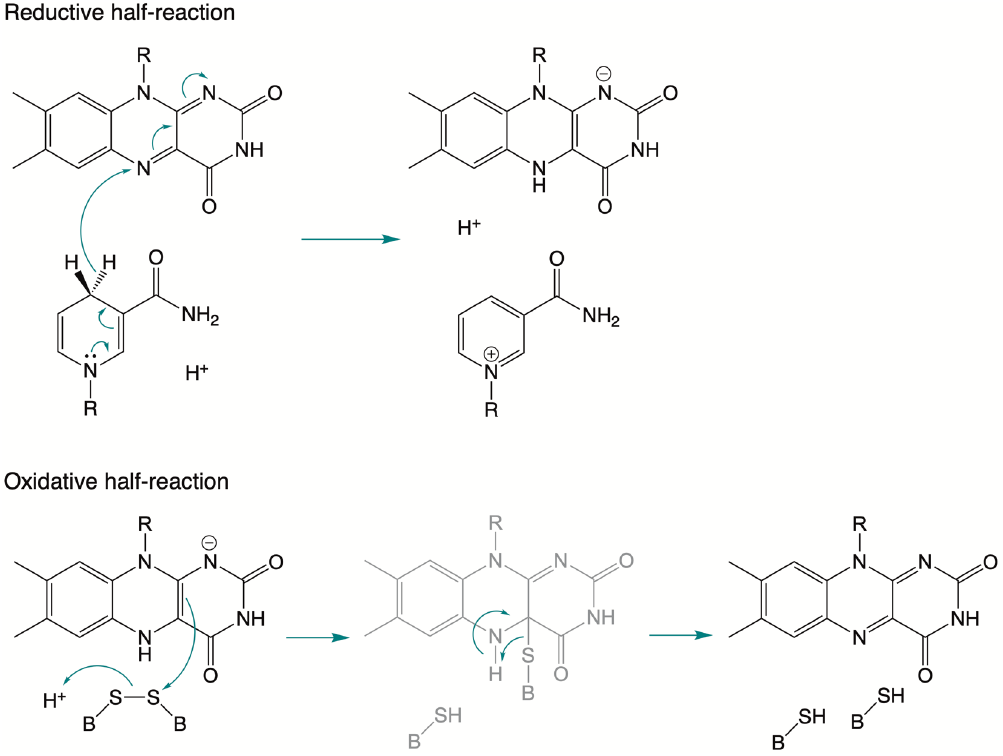
Putative YpdA reaction mechanism. Proposed mechanism for the reduction of BSSB to 2 x BSH. NADPH binds and reduces FAD, NADP^+^ leaves and BSSB binds, and BSSB is reduced to 2 x BSH through a thiol-thiolate pair FAD C4a-cysteine adduct intermediate.

## References

(1) Van Laer, K.; Hamilton, C. J.; Messens, J. Low-Molecular-Weight Thiols in Thiol-Disulfide Exchange. Antioxid. Redox Signal. 2013, 18, 1642–1653.

(2) Chandrangsu, P.; Loi, V. V.; Antelmann, H.; Helmann, J. D. The Role of Bacillithiol in Gram-Positive Firmicutes. Antioxid. Redox Signal. 2018, 28, 445–462.

(3) Couto, N.; Wood, J.; Barber, J. The Role of Glutathione Reductase and Related Enzymes on Cellular Redox Homoeostasis Network. Free Radic. Biol. Med. 2016, 95, 27–42.

(4) Newton, G. L.; Av-Gay, Y.; Fahey, R. C. A Novel Mycothiol-Dependent Detoxification Pathway in Mycobacteria Involving Mycothiol S-Conjugate Amidase. Biochemistry 2000, 39, 10739–10746.

(5) Newton, G. L.; Rawat, M.; La Clair, J. J.; Jothivasan, V. K.; Budiarto, T.; Hamilton, C. J.; Claiborne, A.; Helmann, J. D.; Fahey, R. C. Bacillithiol Is an Antioxidant Thiol Produced in Bacilli. Nat. Chem. Biol. 2009, 5, 625–627.

(6) Sharma, S. V.; Jothivasan, V. K.; Newton, G. L.; Upton, H.; Wakabayashi, J. I.; Kane, M. G.; Roberts, A. A.; Rawat, M.; La Clair, J. J.; Hamilton, C. J. Chemical and Chemoenzymatic Syntheses of Bacillithiol: A Unique Low-Molecular-Weight Thiol Amongst Low G+C GramPositive Bacteria. Angew. Chem. Int. Ed. 2011, 50, 7101–7104.

(7) Hiras, J.; Sharma, S. V.; Raman, V.; Tinson, R. A. J.; Arbach, M.; Rodrigues, D. F.; Norambuena, J.; Hamilton, C. J.; Hanson, T. E. Physiological Studies of Chlorobiaceae Suggest That Bacillithiol Derivatives Are the Most Widespread Thiols in Bacteria. mBio 2018, 9, e01603–18.

(8) Imber, M.; Pietrzyk-Brzezinska, A. J.; Antelmann, H. Redox Regulation by Reversible Protein S-Thiolation in Gram-Positive Bacteria. Redox Biol. 2019, 20, 130–145.

(9) Fowler, V. G.; Miro, J. M.; Hoen, B.; Cabell, C. H.; Abrutyn, E.; Rubinstein, E.; Corey, G. R.; Spelman, D.; Bradley, S. F.; Barsic, B.; Pappas, P. A.; Anstrom, K. J.; Wray, D.; Fortes, C. Q.; Anguera, I.; Athan, E.; Jones, P.; van der Meer, J. T. M.; Elliott, T. S. J.; Levine, D. P.; Bayer, A. S. *Staphylococcus aureus* Endocarditis-a Consequence of Medical Progress. JAMA – J. Am. Med. Assoc. 2005, 293, 3012–3021.

(10) Klevens, R. M.; Morrison, M. A.; Nadle, J.; Petit, S.; Gershman, K.; Ray, S.; Harrison, L. H.; Lynfield, R.; Dumyati, G.; Townes, J. M.; Craig, A. S.; Zell, E. R.; Fosheim, G. E.; McDougal, L. K.; Carey, R. B.; Fridkin, S. K. Invasive Methicillin-Resistant *Staphylococcus aureus* Infections in the United States. JAMA – J. Am. Med. Assoc. 2007, 298, 1763–1771.

(11) Lowy, F. D. Medical Progress – *Staphylococcus aureus* Infections. N. Engl. J. Med. 1998, 339, 520–532.

(12) Livermore, D. M. Antibiotic Resistance in Staphylococci. Int. J. Antimicrob. Agents 2000, 16, S3–S10.

(13) Imber, M.; Loi, V. V.; Reznikov, S.; Fritsch, V. N.; Pietrzyk-Brzezinska, A. J.; Prehn, J.; Hamilton, C.; Wahl, M. C.; Bronowska, A. K.; Antelmann, H. The Aldehyde Dehydrogenase Alda Contributes to the Hypochlorite Defense and Is Redox-Controlled by Protein S-Bacillithiolation in *Staphylococcus aureus*. Redox Biol. 2018, 15, 557–568.

(14) Ghezzi, P. Protein Glutathionylation in Health and Disease. Biochim. Biophys. Acta-Gen. Subj. 2013, 1830, 3165–3172.

(15) Loi, V. V.; Rossius, M.; Antelmann, H. Redox Regulation by Reversible Protein S-Thiolation in Bacteria. Front. Microbiol. 2015, 6, 187.

(16) Lee, J. W.; Soonsanga, S.; Helmann, J. D. A Complex Thiolate Switch Regulates the *Bacillus subtilis* Organic Peroxide Sensor OhrR. Proc. Natl. Acad. Sci. U. S. A. 2007, 104, 8743–8748.

(17) Chi, B. K.; Gronau, K.; Mader, U.; Hessling, B.; Becher, D.; Antelmann, H. S-Bacillithiolation Protects against Hypochlorite Stress in *Bacillus subtilis* as Revealed by Transcriptomics and Redox Proteomics. Mol. Cell. Proteomics 2011, 10, M111.009506.

(18) Chi, B. K.; Roberts, A. A.; Huyen, T. T. T.; Basell, K.; Becher, D.; Albrecht, D.; Hamilton, C. J.; Antelmann, H. S-Bacillithiolation Protects Conserved and Essential Proteins against Hypochlorite Stress in Firmicutes Bacteria. Antioxid. Redox Signal. 2013, 18, 1273–1295.

(19) Gaballa, A.; Chi, B. K.; Roberts, A. A.; Becher, D.; Hamilton, C. J.; Antelmann, H.; Helmann, J. D. Redox Regulation in *Bacillus subtilis:* The Bacilliredoxins BrxA(YphP) and BrxB(YqiW) Function in De-Bacillithiolation of S-Bacillithiolated OhrR and MetE. Antioxid. Redox Signal. 2014, 21, 357–367.

(20) Mikheyeva, I. V.; Thomas, J. M.; Kolar, S. L.; Corvaglia, A. R.; Gaiotaa, N.; Leo, S.; Francois, P.; Liu, G. Y.; Rawat, M.; Cheung, A. L. YpdA, a Putative Bacillithiol Disulfide Reductase, Contributes to Cellular Redox Homeostasis and Virulence in *Staphylococcus aureus*. Mol. Microbiol. 2019, 11, 1039–1056.

(21) Linzner, N.; Loi, V. V.; Fritsch, V. N.; Tung, Q. N.; Stenzel, S.; Wirtz, M.; Hell, R.; Hamilton, C. J.; Tedin, K.; Fulde, M.; Antelmann, H. *Staphylococcus aureus* Uses the Bacilliredoxin (BrxAB)/Bacillithiol Disulfide Reductase (YpdA) Redox Pathway to Defend against Oxidative Stress under Infections. Front. Microbiol. 2019, 10, 1355.

(22) Lofstad, M.; Gudim, I.; Hammerstad, M.; Røhr, Å. K.; Hersleth, H.-P. Activation of the Class Ib Ribonucleotide Reductase by a Flavodoxin Reductase in *Bacillus cereus*. Biochemistry 2016, 55, 4998–5001.

(23) Gudim, I.; Hammerstad, M.; Lofstad, M.; Hersleth, H.-P. The Characterization of Different Flavodoxin Reductase-Flavodoxin (FNR-Fld) Interactions Reveals an Efficient FNR-Fld Redox Pair and Identifies a Novel FNR Subclass. Biochemistry 2018, 57, 5427–5436.

(24) Ojha, S.; Meng, E. C.; Babbitt, P. C. Evolution of Function in the “Two Dinucleotide Binding Domains’” Flavoproteins. PLoS Comput. Biol. 2007, 3, 1268–1280.

(25) Bragg, P. D.; Glavas, N. A.; Hou, C. Mutation of Conserved Residues in the NADP(H)-Binding Domain of the Proton Translocating Pyridine Nucleotide Transhydrogenase of *Escherichia coli*. Arch. Biochem. Biophys. 1997, 338, 57–66.

(26) Ho, B. K.; Gruswitz, F. HOLLOW: Generating Accurate Representations of Channel and Interior Surfaces in Molecular Structures. BMC Struct. Biol. 2008, 8, 49.

(27) Lennon, B. W.; Williams, C. H. Reductive Half-Reaction of Thioredoxin Reductase from *Escherichia coli*. Biochemistry 1997, 36, 9464–9477.

(28) Fagan, R. L.; Palfey, B. A., Flavin-Dependent Enzymes; Elsevier: Oxford, U.K., 2010; Vol. 7.

(29) Sharma, S. V.; Arbach, M.; Roberts, A. A.; Macdonald, C. J.; Groom, M.; Hamilton, C. J. Biophysical Features of Bacillithiol, the Glutathione Surrogate of *Bacillus subtilis* and Other Firmicutes. Chembiochem 2013, 14, 2160–2168.

