## Supporting information for "The Crystal Structures of Bacillithiol Disulfide Reductase YpdA Reveal Structural and Functional Insight into a New Type of FAD-Containing NADPH-Dependent Oxidoreductases"

### Section S1. Expression, Purification and Characterization

#### *Expression and Purification of Bc YpdA, Sa YpdA, and Sa YpdA G10A mutant*

pET-22b(+) plasmids containing the genes for *Bc* YpdA (BC1495, *Bc* ATCC 14579, restriction enzymes NdeI and BamHI)[1], *Sa* YpdA (SACOL1520, *Sa* COL, restriction enzymes NdeI and HindIII) or *Sa* YpdA G10A (SACOL1520, *Sa* COL, restriction enzymes NdeI and HindIII) (GenScript) were transformed into competent *Escherichia coli* One Shot™ BL21 (DE3) cells (Invitrogen, Thermo Fischer Scientific). Cells containing either of the three plasmids were grown in Terrific Broth medium containing 100 μg/mL ampicillin. Protein expression was induced by adding isopropyl β-D-thiogalactoside (IPTG) to a final concentration of 0.5 mM at $OD_{600nm}$ = 0.7-0.9, and the cultures were incubated for 12-16 hours at 20°C with vigorous shaking before cells were harvested and frozen at - 20°C. Cells were thawed and dissolved in 100 mM Tris-HCl, pH 7.5, 1 mM DTT, 5 μg/mL DNase, cOmplete Protease Inhibitor Cocktail (Roche) in a 1:4 cell wet weight to buffer ratio and lysed by sonication. Alternatively, *Sa* YpdA, as well as *Sa* YpdA G10A, were dissolved in 50 mM Tris-HCl, pH 7.5, 500 mM NaCl, 5 μg/mL DNase, cOmplete Protease Inhibitor Cocktail (Roche), prior to cell lysis. The suspensions were centrifuged at 48,000g and the lysates were cleared from nucleic acids by streptomycin sulfate (2.5 %) precipitation, followed by centrifugation at 48,000g. From the lysates, these Tag-free proteins were precipitated with ammonium sulfate ($(NH_4)_2SO_4$) to final concentrations of 0.25 g/mL and 0.22 g/mL for *Bc* and *Sa* YpdAs, respectively, and centrifuged at 39,000g. Proteins were dissolved in 50 mM Tris-HCl, pH 7.5, 1 mM DTT, and desalted using a HiTrap Desalting column (GE Healthcare). Desalted proteins were applied to a HiTrap HP Q column and eluted with linear or step-wise 0-0.5 M KCl or NaCl gradients. As a final polishing step, protein used for crystallization experiments were purified on a Superdex 200 or Superdex 200 Increase column (GE Healthcare) in 50 mM Hepes, pH 7.5, 100 mM KCl. All chromatographic steps were performed using an Äkta purifier FPLC system (GE Healthcare). Protein fractions were pooled, concentrated in Amicon Ultra-15 filter units (10 or 30 kDA MWCO, Merck-Millipore), flash-frozen in liq $N_2$, and stored at - 80°C.

#### *Preparation of Se-Methionine Derivatives*

The L-selenomethionine (Se-Met) derivatives of *Bc* and *Sa* YpdA were expressed and purified in a manner similar to that of the wild type proteins, with the following modifications. Cells were grown at 37°C in M9 minimal medium supplemented with L-methionine (50 mg/L) until the $OD_{600nm}$ had reached 1, harvested, and resuspended in fresh M9 minimal medium without methionine. Cells were further incubated at 37°C until the addition of lysine, phenylalanine, and threonine (100 mg/L of each); isoleucine, leucine, and valine (50 mg/L of each); and L-selenomethionine (50 mg/L), prior to induction with 0.5 mM IPTG (at $OD_{600nm}$ = 1) and incubation for 16 hours at 20°C with vigorous shaking before harvesting and freezing of the cell paste. Cell lysis and protein purification was performed as described for the wild type proteins.

#### *Dynamic Light Scattering (DLS) and Native Polyacrylamide Gel Electrophoresis (PAGE) analyses of protein oligomerization*

In order to estimate the size and molecular weight of the purified *Sa* and *Bc* YpdA proteins, oligomeric states of the YpdA proteins were investigated by DLS (Zetasizer Nano), in 50 mM Hepes, pH 7.5, 100 mM KCl, at 25°C and with 50 μM protein concentrations, in three replicates each. In addition, protein samples of both *Sa* and *Bc* YpdA were analyzed on Native PAGE (NativePAGE™ Novex® 4-16% Bis-Tris Protein Gels, Thermo Fisher Scientific).

#### *UV-vis Spectroscopy of Native and Mutant YpdAs*

UV-vis absorption spectroscopy was performed on 10 μM purified *Sa* YpdA, *Bc* YpdA, and *Sa* YpdA G10A mutant proteins (Agilent Cary 60) to examine the flavin-bound states of the native YpdAs and the apo-form of the G10A mutant. Concentrations were assessed from the absorbance at 453 nm ($\varepsilon_{453nm}$ = 11.5 $mM^{-1}cm^{-1}$) [1] (native proteins) and 280 nm ($\varepsilon_{280nm}$ = 32.3 $mM^{-1}cm^{-1}$) (mutant protein).

### Section S2. Activity measurements

#### *Preparation of BSSB substrate*

As bacillithiol disulfide (BSSB) is not commercially available, an oxidation of reduced BSH was performed, as described by Hamilton and coworkers [2]. In short, prior to enzymatic assays, a solution of $NH_4HCO_3$ solution was added to reduced BSH (Jema Biosciences) dissolved in water at room temperature and stirred with exposure to air for 1 h, flash-frozen in liq $N_2$, and stored at - 80°C.

#### *Enzymatic Assays of BSSB Reduction by Sa and Bc YpdAs*

The reduction of BSSB by NADPH-dependent YpdAs was investigated through spectroscopic determination of the reductase activity. Activity was verified through following the decrease in absorption caused by the oxidation of NADPH at 340 nm. Due to the oxygen sensitivity seen for both *Sa* and *Bc* YpdAs, from the consumption of NADPH regardless of substrate present, enzymatic assays were performed under anaerobic conditions in a glovebox (Plas-Labs Anaerobic Chamber 855-AC and an Agilent 8453 diode-array UV-vis spectrophotometer). The *Sa* YpdA G10A mutant was used as a control, proving the importance of Gly10 in binding of the FAD cofactor, crucial for enzymatic activity. All solutions were degassed on a Schlenk line before transfer to the glovebox. Buffers and stock solutions were sparged with argon for minimum 2 hours in vented vials, and protein samples were subjected to 5-6 cycles of evacuation and refilling with argon. Assays were performed at 25°C in 20 mM Tris, 1.25 mM EDTA, pH 8.0, 240 μM NADPH, with the addition of 12.5 μM purified *Sa* YpdA, *Bc* YpdA or *Sa* YpdA G10A mutant, followed by the addition of 50 μM BSSB, and spectra were collected every second for 250 seconds at 340 nm. Prior to addition of BSSB, reactions were run for 2 min to allow for the reduction of trace amounts of dioxygen by YpdA, in the presence of NADPH. The initial enzymatic assays performed in the presence of dioxygen were executed at 25°C in 20 mM Tris, 1.25 mM EDTA, pH 8.0, 500 μM NADPH, and 50 μM BSSB, followed by the addition of 12.5 μM purified YpdA after 1 min, and spectra were collected every second for 360 seconds at 340 nm (Agilent Cary 60). All assays were performed in replicates.

### Section S3. Protein crystallography

#### *Protein Crystallization*

All initial crystallization screening was performed with a Mosquito crystallization robot (SPT Labtech). Conditions that identified initial hits were further optimized by systematic optimization using the sitting drop vapor diffusion method. Native *Sa* YpdA crystals (57 mg/mL) were obtained with condition A7 from the Morpheus crystallization screen (Molecular Dimensions) (0.03 M magnesium chloride, 0.03 M calcium chloride, 0.1 M MOPS/HEPES-Na, pH = 7.5, 10% w/v PEG 4000, 20% v/v glycerol). Native *Bc* YpdA crystals (22 mg/mL) were obtained with condition B8 from the JCSG-*plus* crystallization screen (Molecular Dimensions)

(0.2 M magnesium chloride hexahydrate, 0.1 M Tris, pH = 7.0, 10% w/v PEG 8000), and briefly soaked in cryoprotectant solution containing mother liquor and 20% ethylene glycol. Diffraction-quality crystals for the Se-Met variant of *Bc* YpdA (25 mg/mL) were grown in condition G3 from the Morpheus crystallization screen (Molecular Dimensions) (0.02 M sodium formate, 0.02 M ammonium acetate, 0.02 M trisodium citrate, 0.02 M sodium potassium L-tartrate, 0.02 M sodium oxamate, 0.1 M MES/imidazole, pH = 6.5, 10% w/v PEG 4000, 20% v/v glycerol). All crystals were grown at room temperature, and flash-frozen in liq $N_2$ prior to data collection.

#### *Crystal Data Collection, Processing, and Refinement*

Diffraction data were collected at MAX IV, Lund, Sweden, on beam line BioMAX (*Sa* YpdA and Se-Met *Bc* YpdA) and at ESRF, Grenoble, France, on beam line ID23-1 (*Bc* YpdA) through MXCuBE3 [3] and ISPyB [4] at 100 K. The Se-Met *Bc* YpdA diffraction data set was collected at 0.976252 Å, a few eV above the theoretical Se absorption K-edge. Diffraction data were indexed and integrated with iMosflm [5] (*Bc* YpdA), auto-processed with EDNA [6] and XDS [7] (*Sa* YpdA), and autoPROC [8] and XDS (Se-Met *Bc* YpdA), and scaled and merged with Aimless in the CCP4 package [9].

Although BLAST sequence alignment searches against the PDB databases indicated hits with up to 30% sequence identity, extended molecular replacement trials with different search models were not successful in solving the structure. This was most likely due to the number of subunits in the asymmetric unit, 4 and 8 combined for *Bc* and *Sa*, respectively, with the possibility of relative movements of the FAD and NADPH binding domains.

The Se-Met *Bc* YpdA structure was solved by single-wavelength anomalous dispersion (SAD) experiment. Matthews coefficient [10] analysis indicated four molecules in the asymmetric unit with a Matthews coefficient of 2.6 $Å^3$/Da and solvent content of 53.2%, indicating that the anomalous signal was significant to about 3.8 Å, so the data set was scaled to 3.5 Å with an anomalous redundancy of 6.6. To solve the structure with SAD, CRANK2 [11] was used through CCP4 online using SFtools, SHELX, SHELXD, REFMAC5, PEAKMAX, MAPRO, Solomon, Multicomb, Parrot, and Buccaneer [12-18]. The solved structure contained four molecules in the asymmetric unit, and initially refined to an R-factor of 36%. This initial model was used as the starting model for the higher resolution native *Bc* YpdA data set of 1.6 Å resolution. After an initial refinement with REFMAC5 further automatic model building was performed with ARP/wARP [19-21] through the CCP4i. This was followed by several cycles of refinement with phenix.refine [22] in the Phenix suite [23] and model building in Coot [24]. Model validation was performed using MolProbity [25]. All structure figures were prepared with PyMOL (Schrödinger, LLC).

The 2.9 Å *Sa* YpdA structure was solved through molecular replacement (MR) using the *Bc* YpdA structure as a starting model. Eight molecules in the asymmetric unit would give a Matthews coefficient of 2.8 $Å^3$/Da and solvent content of 55.6%. To solve the full structure with eight molecules in the asymmetric unit with Phaser [26] through CCP4i, one had to search for one copy of the *Bc* YpdA tetramer, and four copies of the *Bc* YpdA monomer. This was followed by several cycles of refinement initially with REMAC5 and subsequently phenix.refine in the Phenix suite, and model building in Coot. Model validation was performed using MolProbity.

For *Bc* YpdA, all residues have been modelled for chains A and B. For chain C, the C-terminal residue has been excluded, and for chain D, the N-terminal and C-terminal residues have been excluded due to poor electron density. It can additionally be noted that some of the modelled loops have limited electron density. For *Sa* YpdA, the five C-terminal residues have not been modelled for any of the subunits, and some of the modelled loops have limited electron density. Chains G and H have large areas with poor electron density, show high temperature factors, and are more distorted than the other chains. Due to the very limited electron density in some areas of chain G, residues 159-215 were not built into the model for chain G.

Clear electron density was observed for FAD and modelled in all subunits of both *Bc* YpdA and *Sa* YpdA, although with less clear density in the two more distorted chains G and H in *Sa* YpdA (Figure S13A,B). No electron density was observed for NADPH in the *Bc* YpdA structure where both conformations were observed for the possible gating residue Tyr133 (Figure 3F,G). *Sa* YpdA showed clear electron density for NADPH in chains C, D and F, which was accompanied with the closed conformation of residues 295-301, and Tyr128 in open conformation with hydrogen bonding to NADPH (Figure 3D,E,F S13B,D,E). NADPH was not observed in chains A, B and E, which was accompanied with the open conformation of residues 295-301, and Tyr128 in the closed conformation (Figure 3D,E,G). In chains C and F, one conformation of NADPH was modelled in, while in chain D, two orientations of NADPH could be observed accompanied by a movement of Phe51 (Figure 3H).

The standard Phenix restraints used for the FAD cofactor were modified for the 1.6 Å resolution structure, to take into account potential X-ray radiation-induced reduction of the FAD cofactor making the isoalloxazine ring free to bend along the N5–N10 axis (butterfly bend) [27]. The angle of the butterfly bend of the isoalloxazine ring was calculated with the psico module in PyMOL, by calculating the angle between the two planes defined by atoms N5, C4X, C4, N3, C2, C10, N10, and N5, C5X, C3, C7, C8, C9, C9A, N10. A slight butterfly bend of average 4.2° was observed for the isoalloxazine rings in *Bc* YpdA, consistent with expected radiation-induced reduction (Figure S13C).

#### *Crystal packing*

The *Bc* YpdA $P22_12_1$ crystals contained a homo tetramer in the asymmetric unit (Figure S4A), with the monomers having an RMSD value of 1.2-1.3 Å relative to each other (Figure S4B). The *Sa* YpdA $P6_122$ crystals contained eight monomers (RMSD values 1.2-2.8 Å relative to each other, Figure S4E) in the asymmetric unit; a tetramer and two dimers (Figure S4C). The *Sa* YpdA tetramer is similar to the *Bc* tetramer as seen from the overlay in Figure S4D (RMSD value of 1.7 Å). The closest symmetry equivalent subunits in the *Sa* YpdA crystal to the two dimers are shown in Figure S4F (yellow and pink), which shows that they also are part of tetramers. Two of the tetramers (Figure S4F, palecyan and yellow) overlay well (RMSD value of 1.3 Å), while for the third tetramer (pink), each of the two dimers overlay well (RMSD value of 2.5 Å) with the two dimers of the other tetramers, however, overlaying one dimer results in the second dimer to be shifted 15 Å relative to the others (Figure S4G). Therefore, the *Sa* YpdA $P6_122$ crystal also contains tetramers, however, most likely due to crystal packing, one of the tetramers had to adopt by slightly sliding the two dimers relative to each other (Figure S4G).

### Section S4. Bioinformatics and Structural Analysis

#### *BLAST search for homologous sequences*

The *Sa* YpdA sequence (Locus tag SACOL1520) was used for a BLAST (Basic Local Alignment Search Tool) search using the NCBI web interface (https://blast.ncbi.nlm.nih.gov/Blast.cgi) searching the reference proteins (refseq_protein) database searching for 20,000 sequences with a threshold E-value of $1e^{-6}$ using the BLOSUM62 matrix. The same search was also performed limiting the search to the different bacterial phyla.

The BLAST search resulted in 3977 bacterial organism hits. The dominating bacterial phylum for *Sa* YpdA homologous sequences was Firmicutes (1813 organism hits) followed by Bacteroidetes (1425), Proteobacteria (527), Deinococcus-Thermus (93), Actinobacteria (71) and Acidobacteria (49). Of these organisms, Proteobacteria use GSH as the main LMW thiol, and Actinobacteria use MSH, while low G+C Firmicutes and Deinococcus-Thermus contain BSH, and Bacteriodetes and Acidobacteria to a large extent contain Me-BSH[28]. Therefore, Proteobacteria and Actinobacteria were not further studied, while three of the top hits from Firmicutes, Deinococcus-Thermus, Bacteroidetes, and Acidobacteria were subjected to multiple sequence alignments and phylogenetic tree analysis through Jalview [29]. The alignments were performed with Clustal Omega [30] and tree analysis with average distances using the BLOSUM62 matrix. Figure S9 shows that the sequences fall into four clades corresponding to the four phylum classes the sequences belong to. The sequence identity compared to the *Sa* YpdA sequence was for the other Firmicutes in Figure S9 ~60%, for Bacteroidetes ~44%, Deinococcus-Termus ~37%, and Acidobacteria ~44%, and all these sequences have been annotated as putative YpdAs in the refseq_protein database. A cysteine in *Sa* YpdA has been suggested to function as an active site residue, involved in the reduction of BSSB [31]. This cysteine is found in the canonical FAD binding GXGXXG motif in Firmicutes, however, in Bacteroidetes, this residue is mainly replaced by isoleucine, in Deinococcus-Thermus valine, and in Acidobacteria threonine (Figure S9). Further studies on the proposed YpdAs from e.g. Bacteroidetes, Deinococcus-Thermus, and Acidobacteria need to be performed to reveal if they are YpdAs or if they have other functions.

The putative YpdA ortologs used in this study were from *Staphylococcus aureus, Bacillus cereus, Bacillus subtilis, Chryseobacterium frigidisoli, Fabibacter pacificus, Algoriella xinjiangensis, Meiothermus luteus, Thermus filiformis, Deinococcus misasensis, Acidobacteria bacterium, Candidatus Sulfotelmatobacter sp.,* and *Granulicella sp.*

#### *Structure comparison - structural alignment search with DALI (Distance-matrix ALIgnment)*

A search for similar structures of the YpdA in the Protein Data Bank (PDB) was performed using the DALI protein structure comparison server using the *Bc* YpdA structure as a search template [32]. The most similar monomer structures were flavoprotein monooxygenases (FPMOs), thioredoxin reductases (TrxRs), thioredoxin-like ferredoxin $NADP^+$ oxidoreductases (FNRs), dihydrolipoyl dehydrogenases (DLDs), and gluthatione reductases (GRs) (Table S2).

The monomer structures of YpdA were compared to the monomer structures FPMO (PDBid 4C5O), TrxR (PDBid 5VT3), FNR (PDBid 3AB1), DLD (PDBid 6CMZ), and GR (PDBid 1GRB) from Table S6, and are compared and overlaid (Figure S6). In addition, the multimeric structures are shown. For TrxR, both the FR and FO (PDBid 1TDF) dimeric states are shown, and for FNR, the two structures of the *Bc* FNRs are shown (PDBid 6GAS and 6GAR). Figures were generated with PyMOL.

Although the NADPH and FAD domains of YpdA are similar to DLD and GR (Figure S2, Table S2), DLD and GR have 100-130 additional C-terminal residues. Therefore, these proteins were not included in the structural sequence alignments generated with DALI (Figure S3). The secondary structure assignments were calculated with DSSP [33-34] and shown below the sequence. The figure was generated in JalView and colored by % identity.

***Phylogenetic analysis of YpdA with other flavin oxidoreductases in selected Firmicutes***

Through several BLAST searches the likely sequences of YpdAs, FNRs, TrxRs, and FPMOs in the selected Firmicutes *Staphylococcus aureus subsp. aureus COL*, *Staphylococcus epidermidis* RP62A, *Bacillus cereus ATCC 14589, Bacillus anthracis str. sterne*, and *Bacillus subtilis subsp.* subtilis str. 168 were found. Multiple sequence alignments were performed with Clustal Omega and phylogenetic tree analysis with average distances using the BLOSUM62 matrix (Figure S6). The figures were generated in JalView and the sequence alignments were colored by % identity. The phylogenetic tree analysis shows that the four oxidoreductase types from the five phyla fall into four clades corresponding to the four oxidoreductases types, and that YpdA forms its own clade, indicating that it constitutes a separate type of flavin oxidoreductases (Figure S6).

The locus tags for the sequences used are shown in parentheses for *Sa* YpdA (SACOL1520), *Se* YpdA (SERP1047), *Bc* YpdA (BC1495), *Ba* YpdA (BAS1405), *Bs* YpdA (BSU22950), *Sa* FNR1 (SACOL2369), *Bc* FNR1 (BC0385), *Ba* FNR1 (BAS0337), *Bc* FNR2 (BC4926), *Ba* FNR2 (BAS4797), *Bs* YumC (BSU32110), *Bs* YcgT (BSU03270), *Sa* TrxR (SACOL0829), *Se* TrxR (SERP0432), *Bc* TrxR (BC5159), *Ba* TrxR (BAS5007), *Bs* TrxR (BSU34790), *Sa* FPMO (SACOL2600), *Se* FPMO (SERP2384), *Bc* FPMO (BC3447), *Ba* FPMO (BAS3253), and *Bs* FPMO (BSU26640).

***Analysis of conserved residues of YpdA with ConSurf***

To evaluate the degree of conservation of residues in YpdA, ConSurf [35-38] was run on the *Bc* YpdA structure. The run was based on the homologue search algorithm HMMER, searching sequences from UniRef90, and multiple sequence alignment with MAFFT. This gave 2266 unique HMMER hits, and ConSurf used a sample of 150 sequences that represented the list of homologues sequences to map the conservation on a 9-bin scale from turquoise (most variable) to maroon (most conserved). The conservation color coding was then mapped onto the *Bc* YpdA crystal structure and figures were generated with PyMOL.

Phylogenetic analysis on YpdA was performed in Jalview on the 150 sequences selected in the ConSurf runs (a few outlier sequences were not included). Clustal Omega was used for sequence alignment, and average distances in the phylogenetic tree were calculated with BLOSUM62.

The selected sequences fall into three clades in the phylogenetic trees showing that the most homologous sequences to *Bc* YpdA is found in other Firmicutes, Bacteroidetes and Acidobacteria (Figure S10).

The surface representation of the ConSurf colored YpdA show that the most conserved part of the surface (white-to-maroon) is between the tetramers, which supports the conclusion of YpdA being a biological tetramer (Figure S11).

The most conserved residues (maroon-to-lightmaroon) are found around the FAD cofactor, around the NADPH binding site, and the residues lining the solvent channel spanning the structure in connection with the FAD cofactor.

#### *Analysis of channels with HOLLOW*

The potential channel for BSSB binding was generated with HOLLOW [39] using a 4 Å probe in cylinder mode between Phe51 and Tyr152(*Bc*)/Tyr147(*Sa*), or Lys170(*Bc*)/Lys165(*Sa*) and Tyr152 . Similarly, the sphere mode was used to look for any small channels around Cys14.

#### *Modelling/estimating the position of BSSB or NADP$^+$ in YpdA*

To show potential BSSB binding sites around the FAD group, BSSB was manually positioned in Coot within the HOLLOW-generated channels and regularized with respect to stereochemical restraints in Coot. To span potential orientations within the whole channel, three molecules of BSSB were fitted. One was positioned with the disulfide bond close to the C4a (C4X, FAD numbering) atom of FAD (5 Å).

#### *Ligand interaction generated with LigPlot$^+$.*

To make schematic diagrams of protein-ligand interactions from the protein structure (PDB files), the program LigPlot$^+$ was used with default parameters to show hydrogen bonds and hydrophobic contacts represented by dashed lines and arcs with spokes radiating toward the ligand atoms they contact, respectively [40-41].

**Supplementary Figures and Tables:**

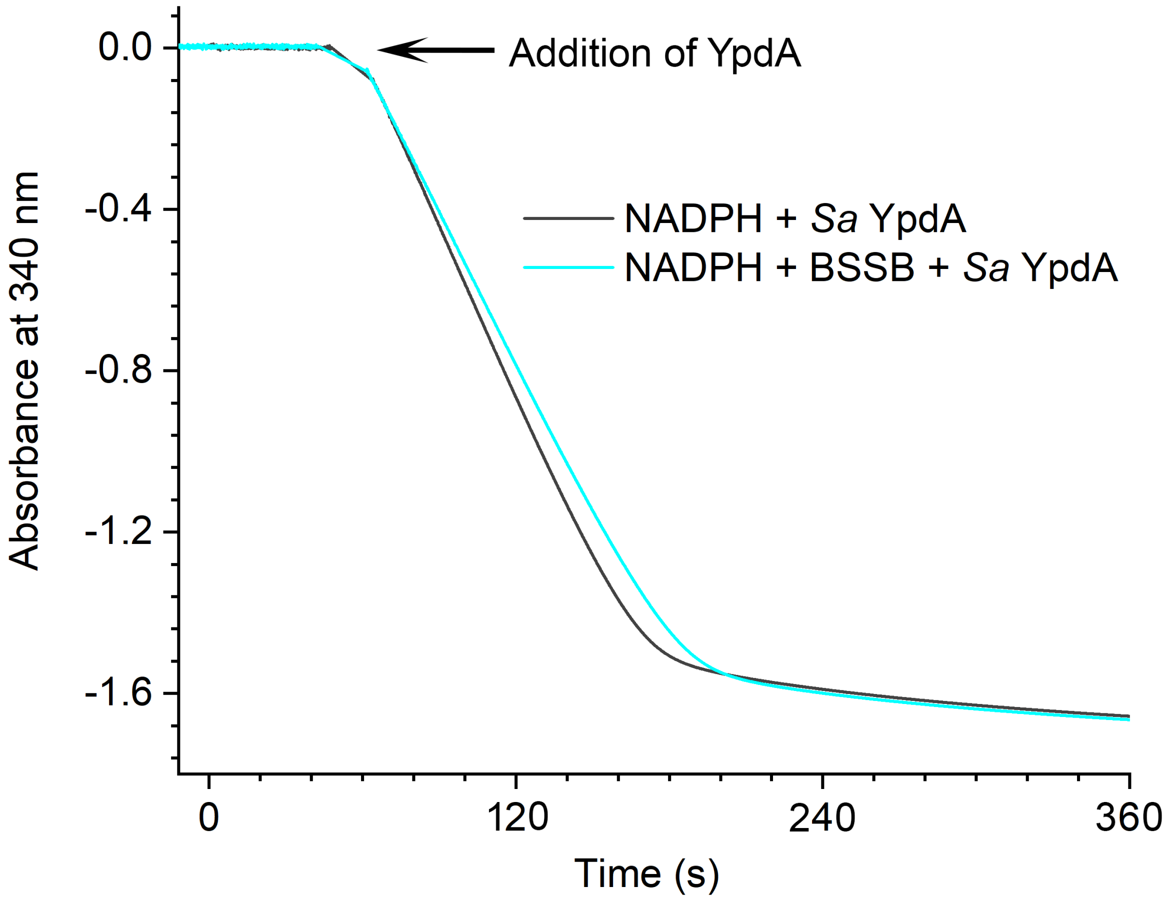

**Figure S1: NADPH consumption by YpdA under aerobic conditions.** YpdA consumes NADPH at similar rates with or without BSSB added to the reaction in the presence of dioxygen.

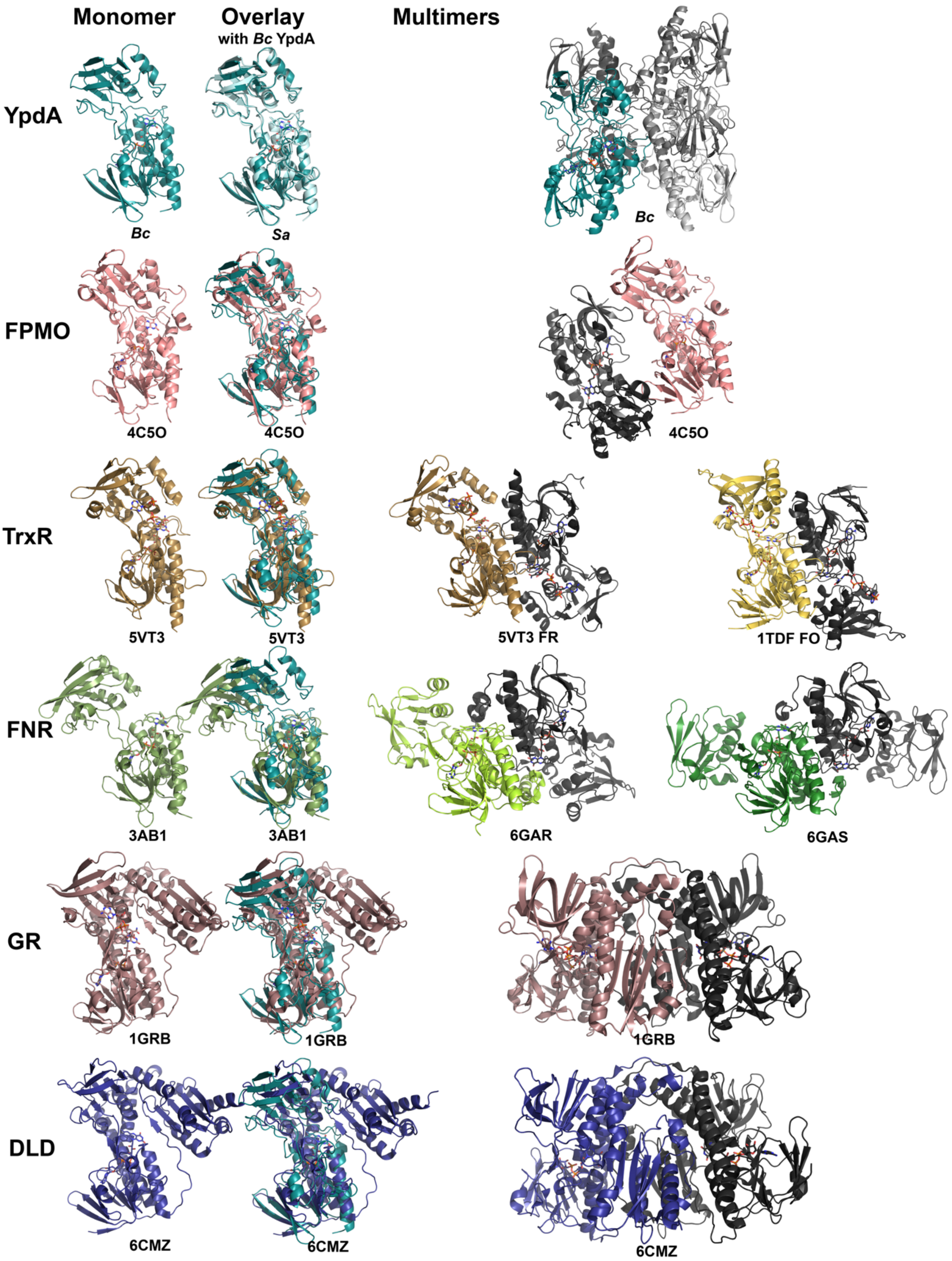

**Figure S2**: **Structure comparison.** Monomer structures of YpdA and top DALI hits for the FPMO, TrxR, FNR, GR, and DLD from Table S2, the monomer structure alignment of FPMO, TxrR, FNR, GR, with YpdA, and selected multimerization structures of the different proteins. The PDBid is listed for each structure.

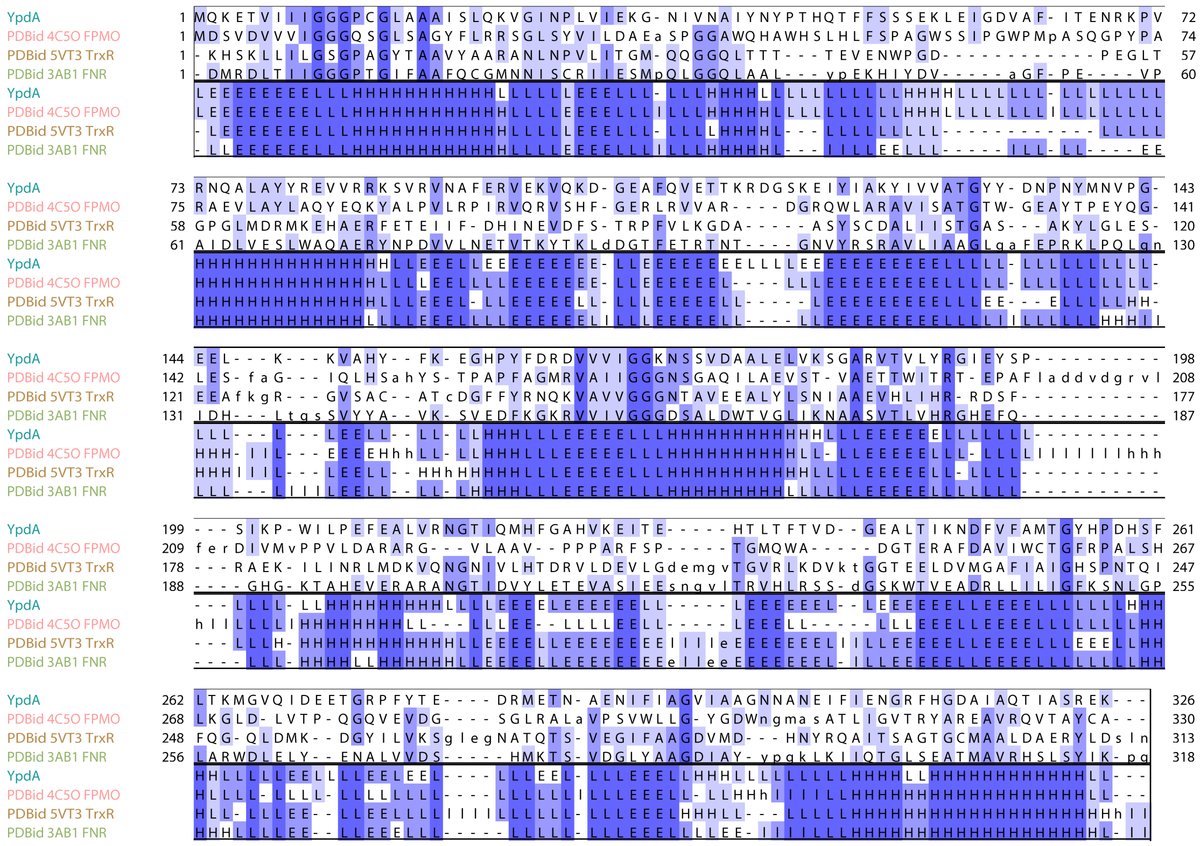

**Figure S3**: **Structural alignment of *Bc* YpdA, FPMO (PDBid 4C5O), TrxR (PDBid 5VT3), and FNR (PDBid 3AB1) performed with a DALI search.** The secondary structure assignments were calculated with DSSP and shown below the sequence. The figure was generated in JalView and colored by % identity.

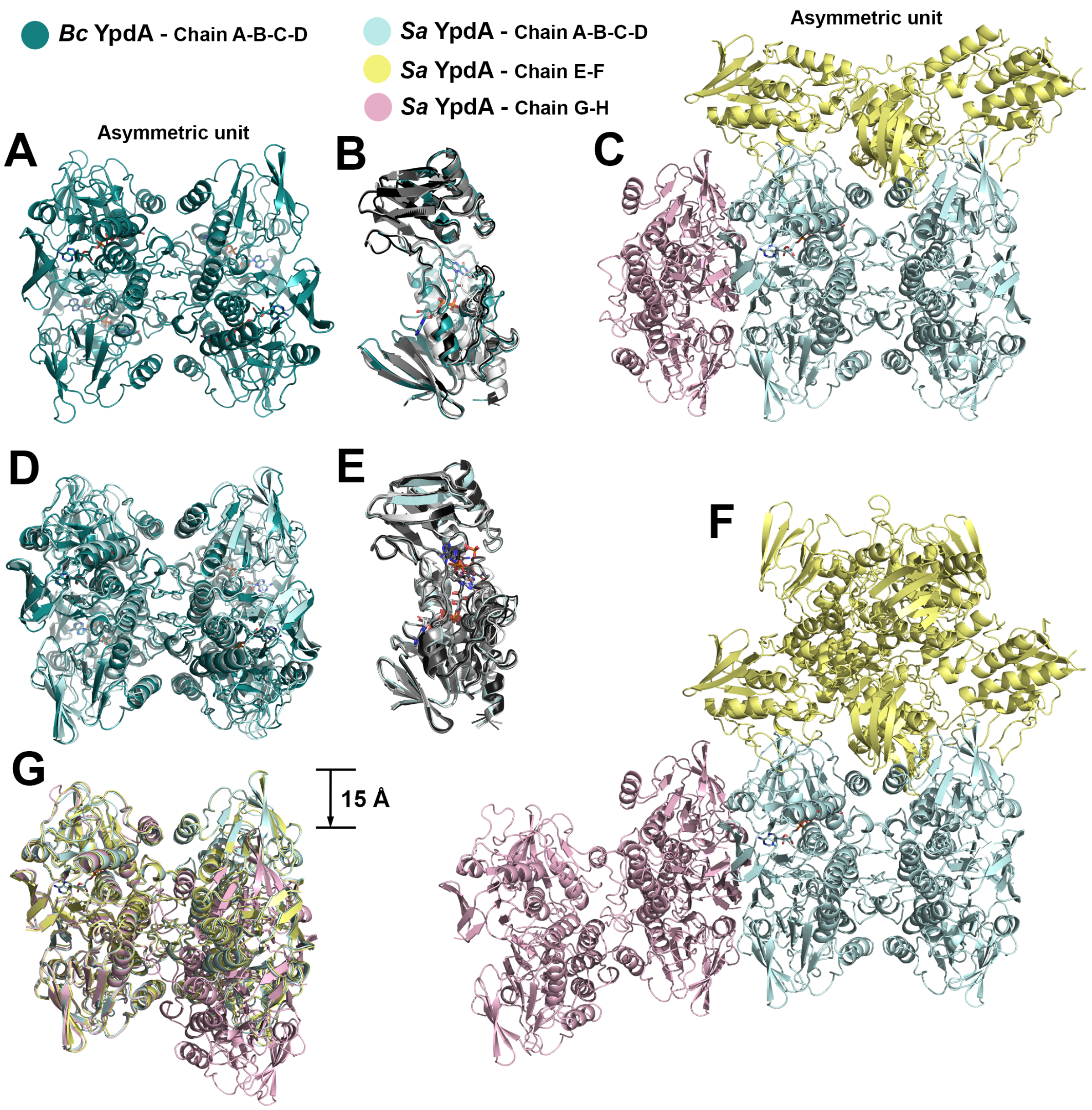

**Figure S4**: **Crystal packing**. (**A**) Asymmetric unit of *Bc* YpdA (tetramer). (**B**) Overlay of the *Bc* YpdA monomers. (**C**) Asymmetric unit of *Sa* YpdA (octamer). (**D**) Overlay of the *Bc* YpdA tetramer with the palecyan colored tetramer of the *Sa* YpdA from B. **(E)** Overlay of the eight *Sa* YpdA monomers from C. (**F)** The octamer from B added the closest dimer symmetry equivalents of the yellow and pink colored dimers of the *Sa* YpdA octamer. (**G)** Overlay the three tetramers shown in F with full overlay of the palecyan and the yellow tetramers. The second dimer of the pink tetramer is shifted 15 Å relative to the first dimer compared, as compared to the palecyan and pink tetramers.

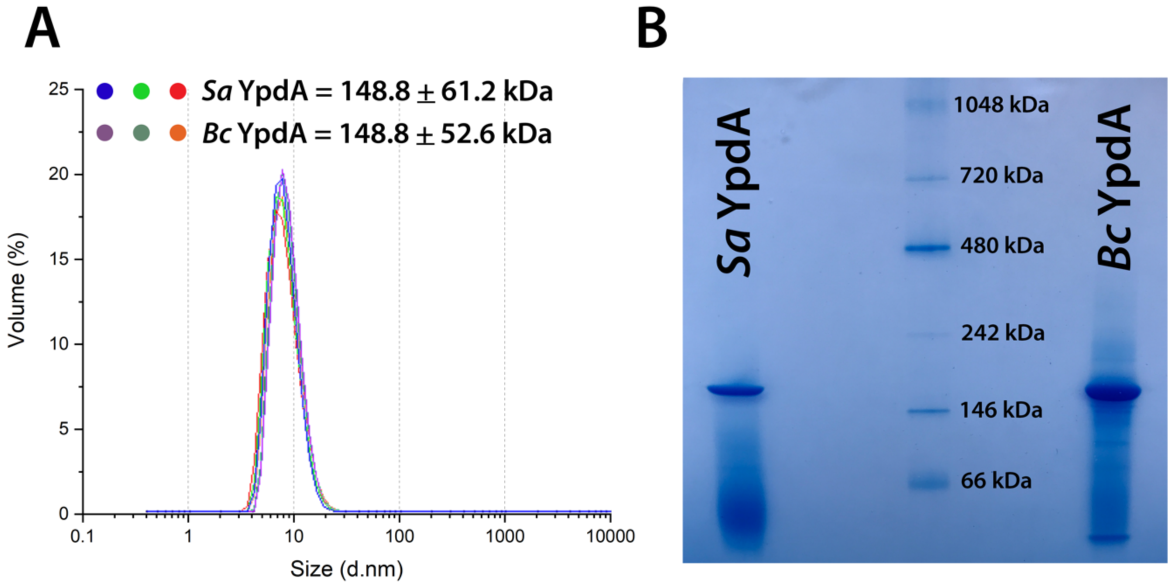

**Figure S5: Molecular weight estimation of YpdA proteins.** Analysis of the oligomerization state of *Sa* YpdA (36.7 kDa/monomer) and *Bc* YpdA (36.5 kDa/monomer) performed with (**A**) DLS, and (**B**) Native PAGE, indicating that YpdA is a biological tetramer.

**A**

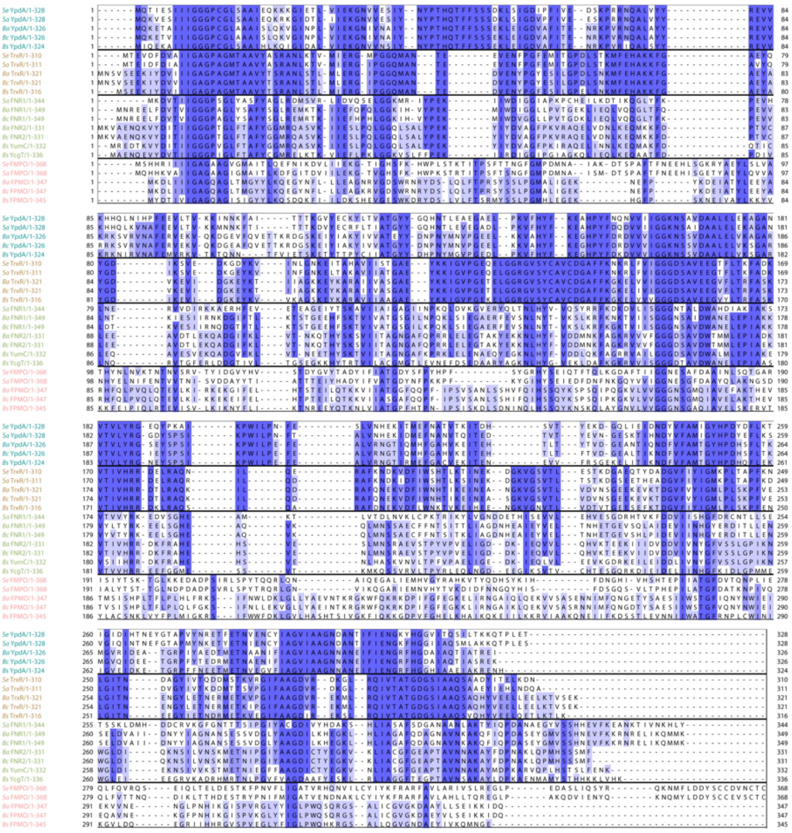

**B**

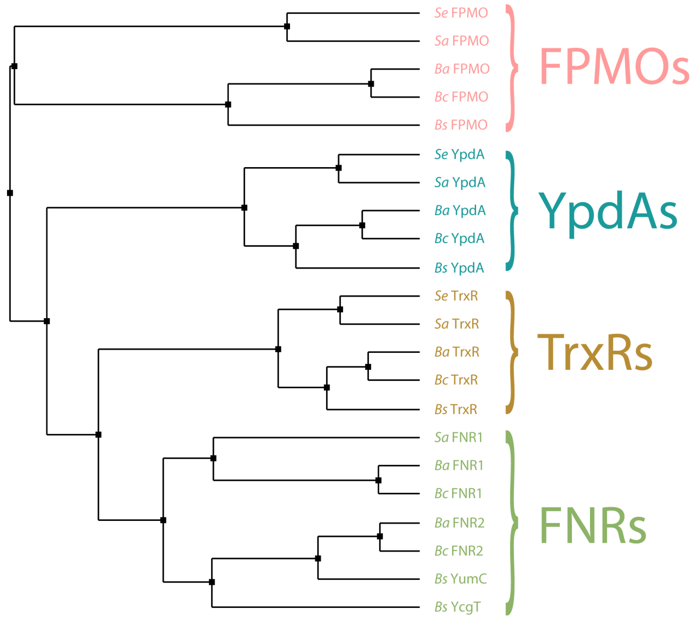

**Figure S6**: **Multiple sequence alignment and phylogenetic tree analysis of proposed YpdAs, FMPOs, FNRs, and TrxRs from the selected Firmicutes *S. aureus*, *S. epidermis*, *B. cereus, B. anthracis*, and *B. subtilis.*** (**A**) Multiple sequence alignment generated with Clustal Omega through Jalview. The coloring is according to % identity. The sequences are grouped according to the phylogenetic tree in B. (**B**) Phylogenetic tree calculated with Jalview with average distances using the BLOSUM62 matrix on the sequence alignment A.

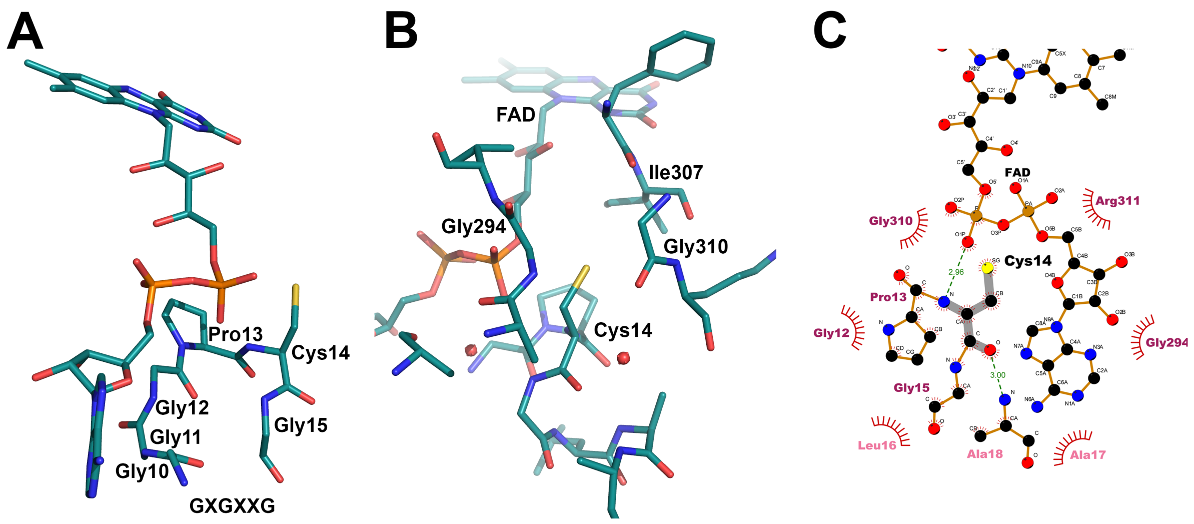

**Figure S7**: **Selected structural features in *Bc* YpdA. (A)** The GGGPCG motif. (**B)** The environment around Cys14. **(C)** LigPlots$^{+}$ showing the interaction between Cys14 and other residues with the residue name colored according to conservation obtained from ConSurf.

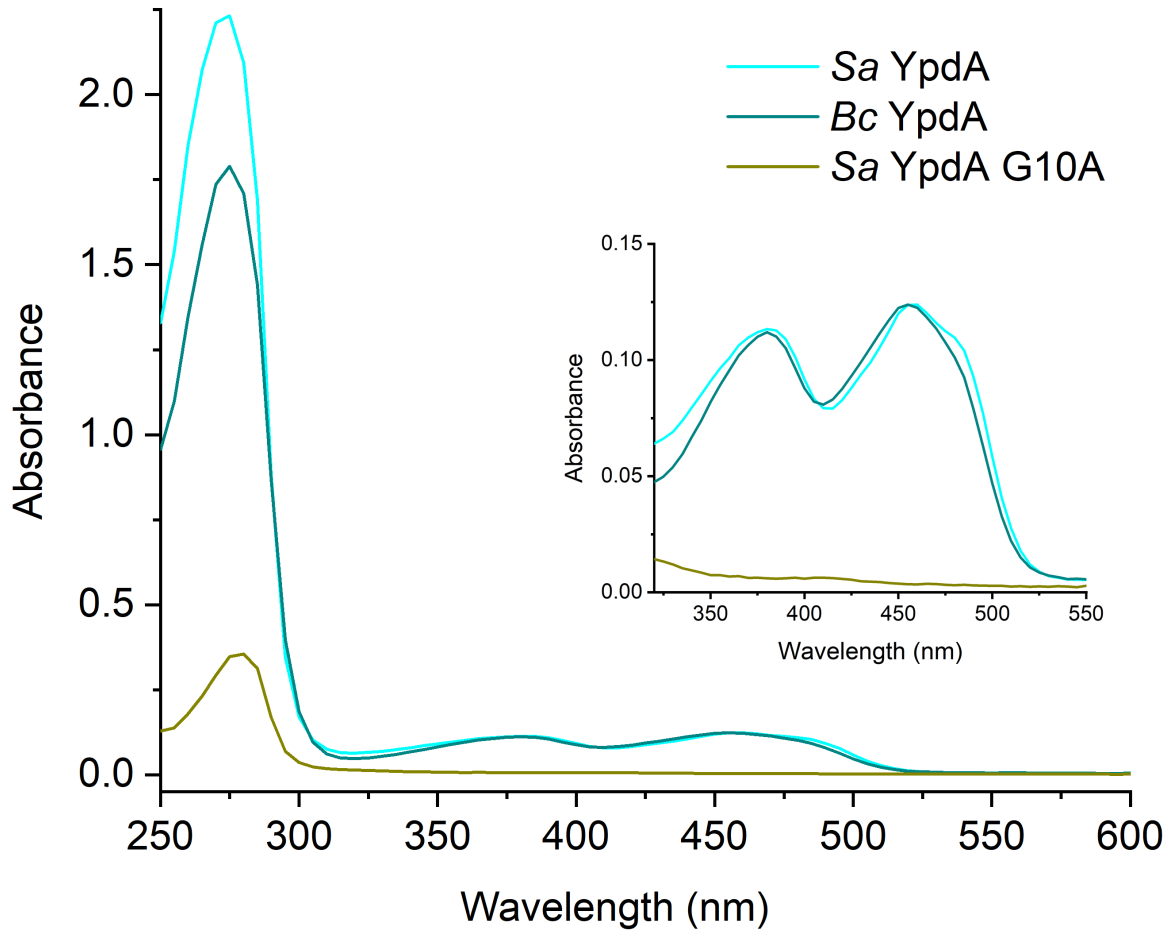

**Figure S8: UV-vis characterization of YpdA proteins.** UV-vis absorption spectra of 10 µM purified *Sa* YpdA, *Bc* YpdA, and *Sa* YpdA G10A mutant. The spectra show typical features of the oxidized state of the FAD cofactor in *Sa* YpdA and *Bc* YpdA (see inset), whereas the YpdA G10A mutant exists in its apo-form.

**A**

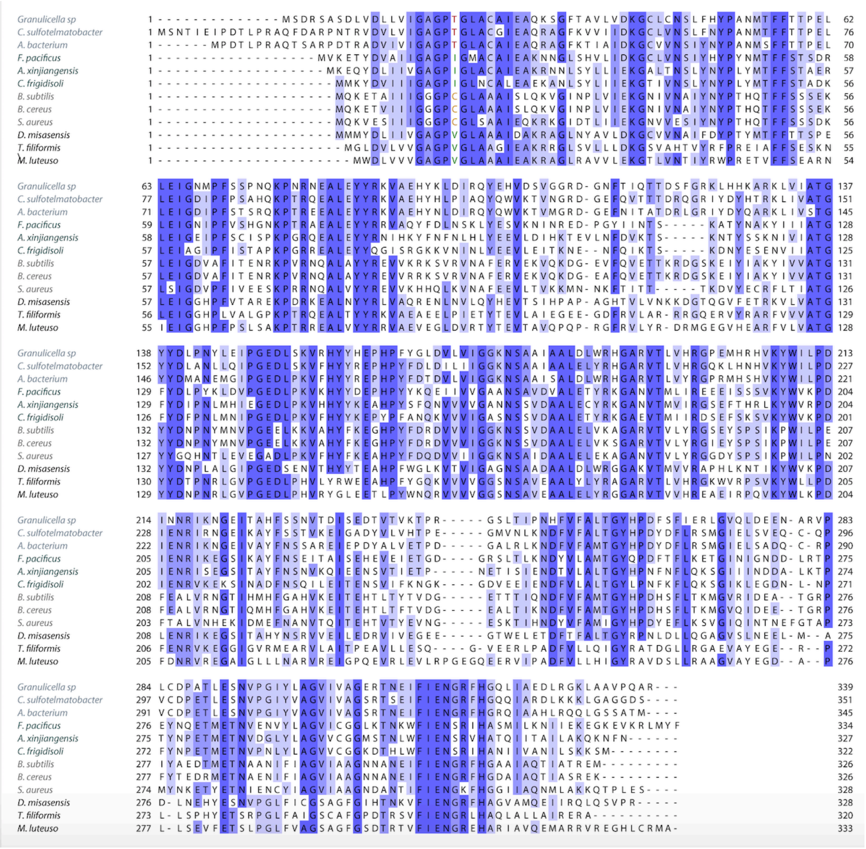

**B**

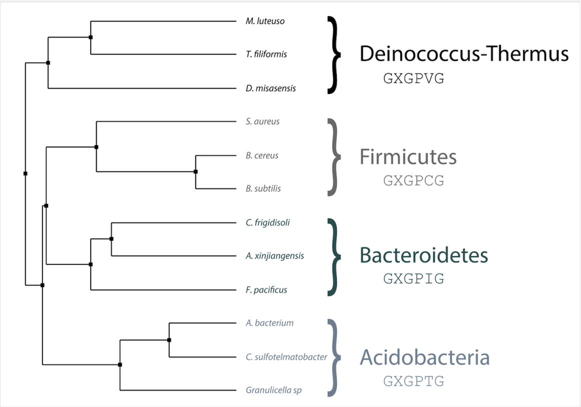

**Figure S9: Multiple sequence alignment and phylogenetic tree analysis.** Analysis of selected YpdA homologous sequences form the phyla Firmicutes, Bacteroidetes, Deinococcus-Thermus and Acidobacteria. (**A**) Multiple sequence alignment generated with Clustal Omega through Jalview. The coloring is according to % identity. The sequences are grouped according to the phylogenetic tree in B. (**B**) Phylogenetic tree calculated with Jalview with average distances using the BLOSUM62 matrix on the sequence alignment in A. The common GXGPXG motif of YpdA is shown for each clade. The putative YpdA orthologs used were from *S. aureus, B. cereus, B. subtilis, C. frigidisoli, F. pacificus, A. xinjiangensis, M. luteus, T. filiformis, D. misasensis, A. bacterium, C. Sulfotelmatobacter sp.,* and *G. sp.*

# A

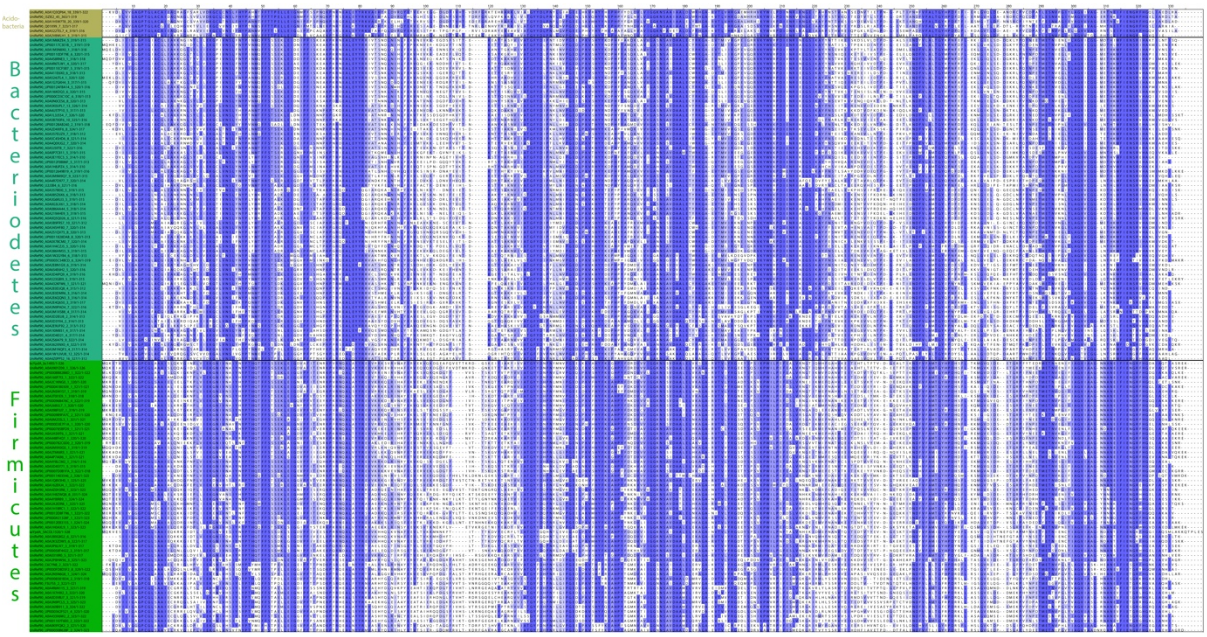

# B

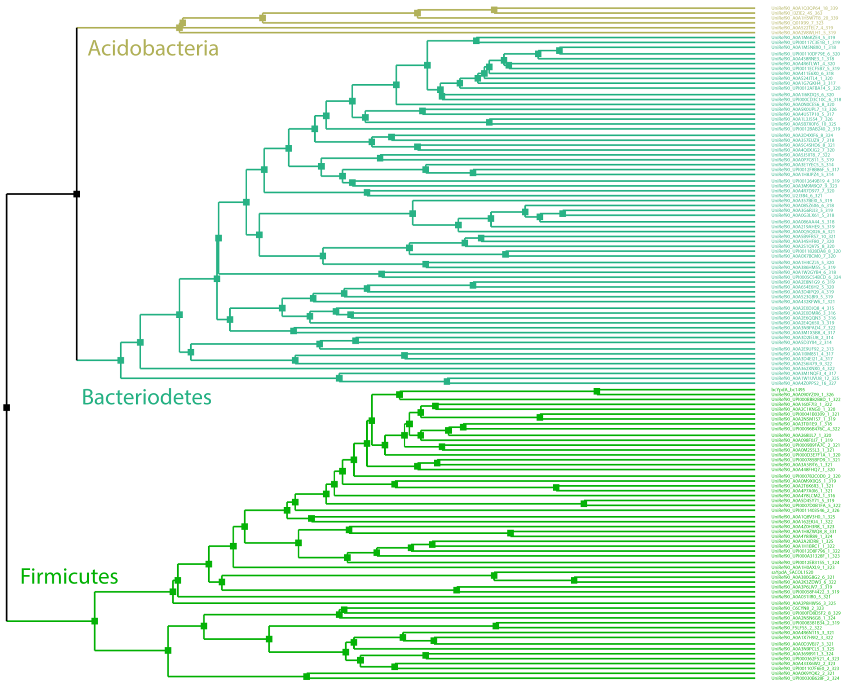

**Figure S10**: **Multiple sequence alignment and phylogenetic tree analysis of the ~150 sampled sequences obtained from a ConSurf search using the *Bc* YpdA structure.** (**A**) Multiple sequence alignment generated with Clustal Omega through Jalview. The coloring is according to % identity. The sequences are grouped according to the phylogenetic tree in B. (**B**) Phylogenetic tree calculated with Jalview with average distances using the BLOSUM62 matrix on the sequence alignment A.

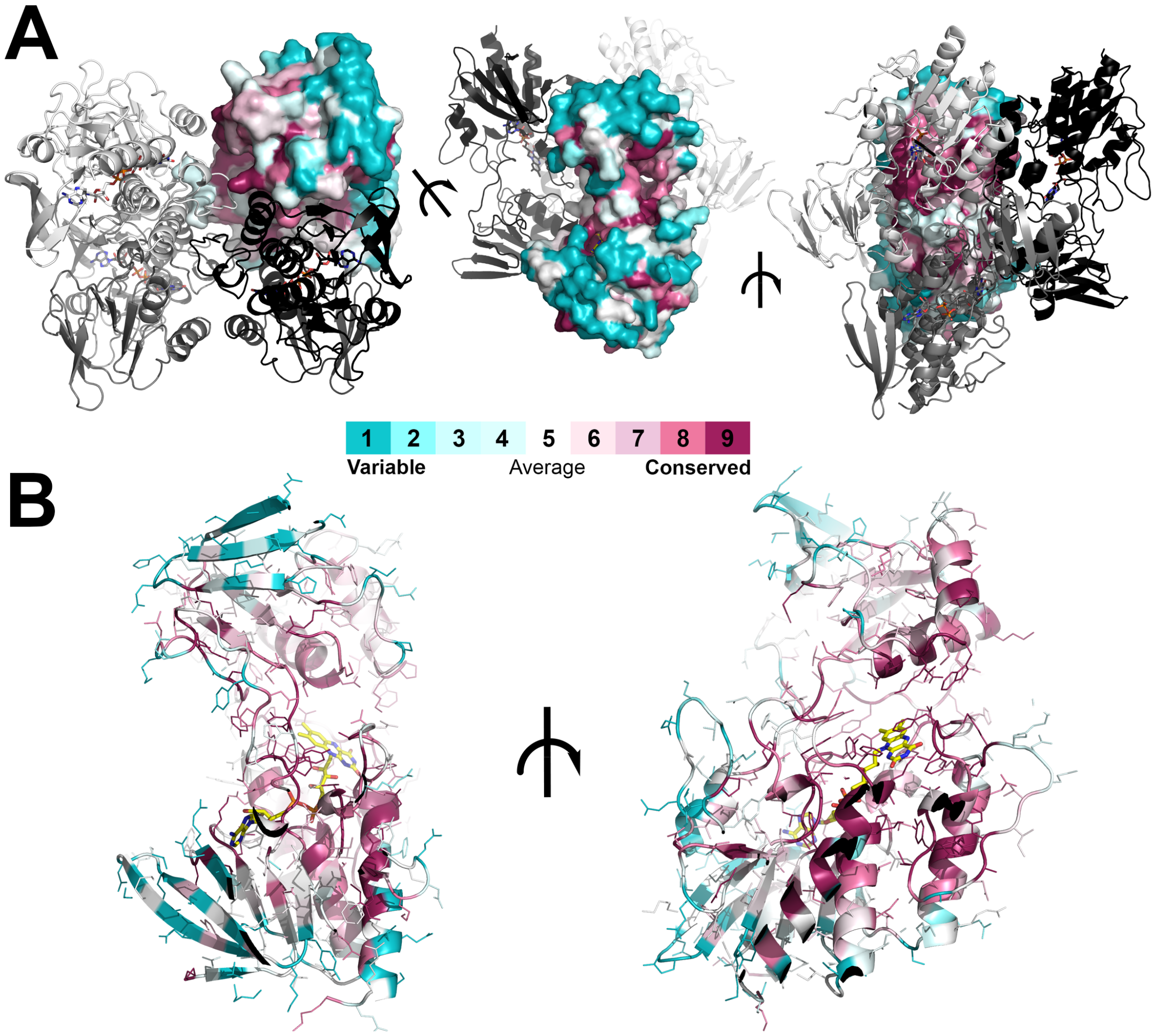

**Figure S11: Conservation of residues in YpdA evaluated using ConSurf.** Variable residues are colored in turquoise, highly conserved residues are colored in maroon, and figures are generated in PyMOL. The FAD cofactor is shown as sticks and colored with yellow carbon atoms. The degree of conservation of the YpdA residues are represented as surface, lines and/or cartoon in the different panels. (**A)** Overall view of the YpdA tetramer in different orientations with the conservation shown for one monomer. (**B)** Monomer view of conserved residues shown in two different orientations.

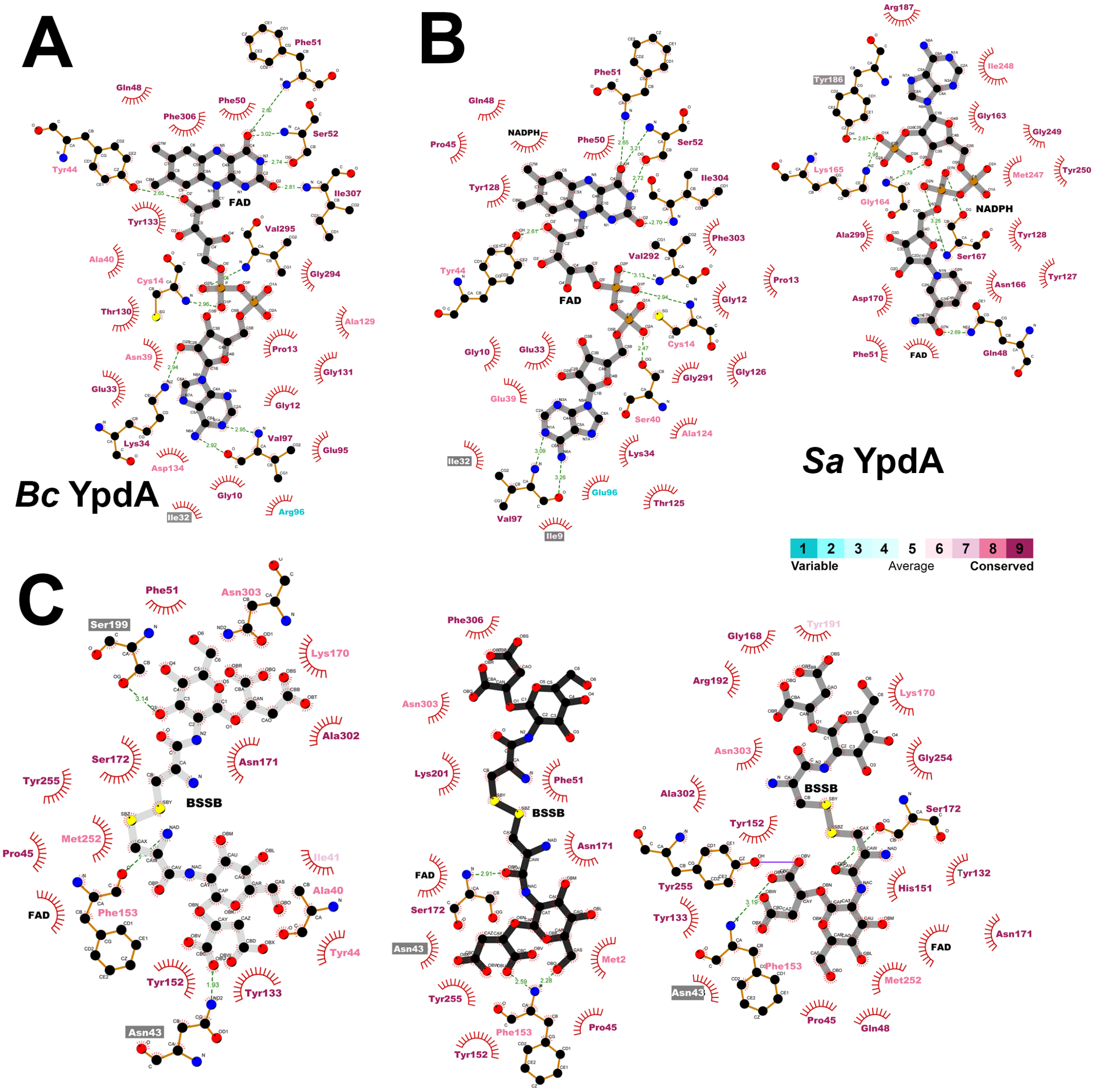

**Figure S12**: **Cofactor and putative BSSB binding sites.** **(A)** LigPlot+ showing the FAD bonding site in *Bc* YpdA and **(B)** FAD and NADPH binding sites in *Sa* YpdA. **(C)** Three LigPlots+ showing the ConSurf colored conserved residues interacting with the three possible BSSB molecules placed into the HOLLOW-generated solvent channel.

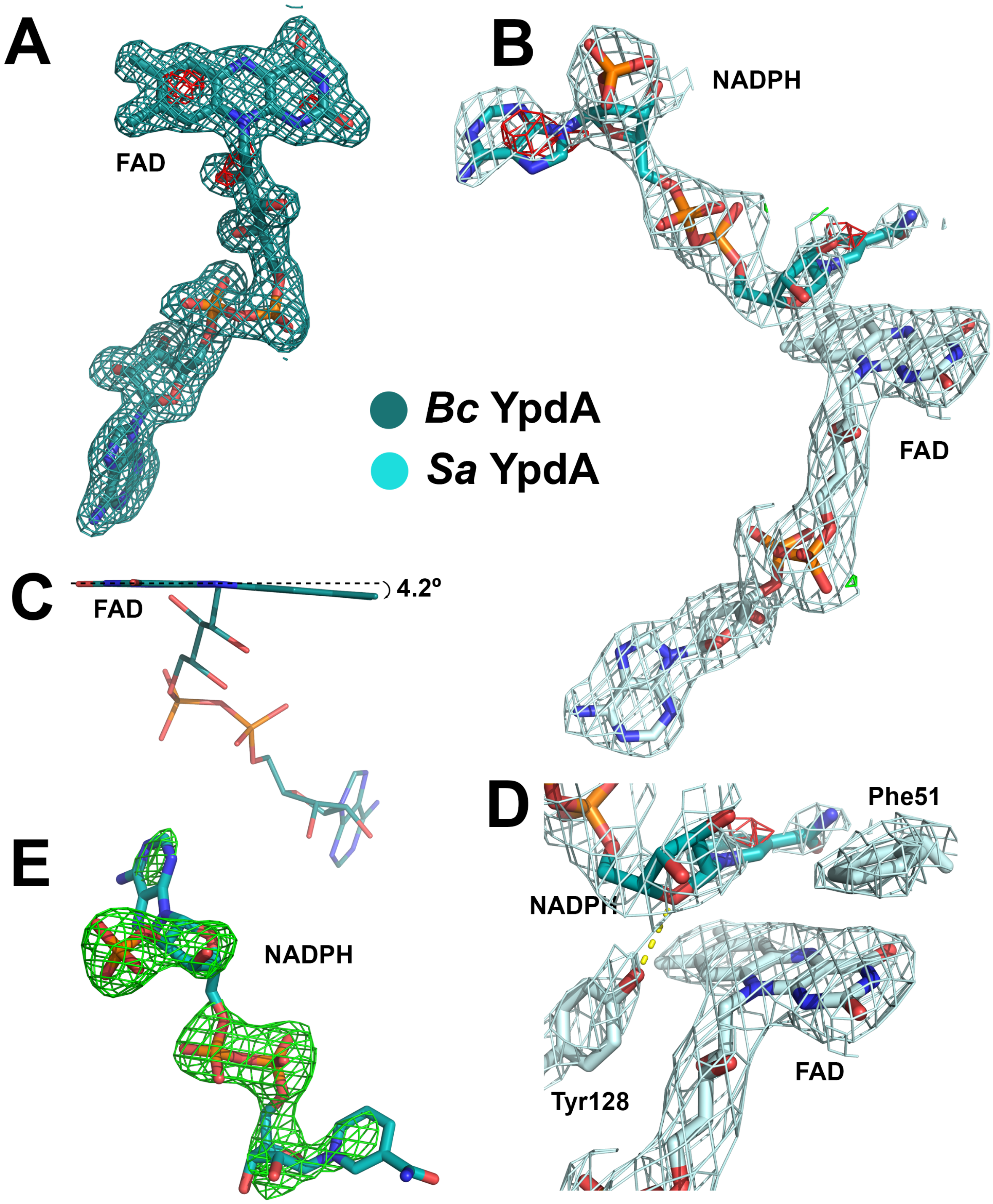

**Figure S13**: **Cofactor electron density, geometry and interactions**. Electron density for the *Bc* YpdA structure chain A **(A)** and *Sa* YpdA structure chain C **(B)**, with the $2F_o$–$F_c$ map contoured at 1σ (colored in darkteal/palecyan) and the $F_o$–$F_c$ maps contoured at ±3 σ (colored in green/red) around the FAD and NADPH cofactors. **(C)** The refined butterfly bending of the flavin plane from likely X-ray radiation induced reduction. **(D)** Electron density for cofactors and residues likely involved in gating of NADPH and possibly BSSB, showing a hydrogen bond from Tyr128 to NADPH in the *Sa* YpdA NADPH-bound state (open conformation). **(E)** Omit electron density map of NADPH in *Sa* YpdA chain C contoured at 3 σ (colored in green).

**Table S1**: **Crystal data collection and refinement statistics.**

| | *Bc* YpdA Se-Met | *Bc* YpdA | *Sa* YpdA |
|---|---|---|---|
| **Data collection** | | | |
| X-ray source | BioMAX, MAXIV | ID23-1, ESRF | BioMAX, MAXIV |
| Detector | EIGER 16M | Pilatus 6M | EIGER 16M |
| Wavelength (Å) | 0.976252 | 0.97242 | 1.49379 |
| Space group | $P22_12_1$ | $P22_12_1$ | $P6_122$ |
| $a$, $b$, $c$ (Å) | 88.8, 115.3, 147.9 | 88.8, 115.0, 146.8 | 179.3, 179.3, 349.3 |
| $\alpha$, $\beta$, $\gamma$ (°) | 90, 90, 90 | 90, 90, 90 | 90, 90, 120 |
| Type of data collection | Standard rotation | Helical scan | Standard rotation |
| Rotation range per image (°) | 0.1 | 0.1 | 0.1 |
| Total rotation range (°) | 360 | 100 | 180 |
| Exposure time per image (s) | 0.011 | 0.037 | 0.011 |
| Flux (ph/s) / Transmission (%) | n/a / 25% | $2.1 \times 10^{12}$ / 100% | n/a / 50% |
| Beam size (μm$^2$) | 50 × 50 | 30 × 45 | 50 × 50 |
| Crystal size (μm$^3$) | 30 × 200 × 200 | 20 × 150 × 250 | 75 × 75 × 250 |
| Mosaicity (°) | 0.26 | 0.17 | 0.08 |
| Resolution range (Å) | 147.9-3.5 (3.8-3.5) | 22.5-1.6 (1.63-1.60) | 49.6-2.9 (2.96-2.90) |
| Total no. of reflections | 254621 | 571284 | 1328276 |
| No. of unique reflections | 19810 | 190500 | 73800 |
| $R_{meas}$ | 0.184 (0.344) | 0.084 (1.036) | 0.169 (0.580) |
| $R_{merge}$ | 0.176 (0.330) | 0.069 (0.860) | 0.165 (0.565) |
| Completeness (%) | 99.8 (99.3) | 96.7 (96.3) | 99.8 (99.4) |
| Anomalous completeness | 99.9 (99.9) | | |
| Multiplicity | 12.9 (13.1) | 3.0 (3.1) | 18.0 (20.4) |
| Anomalous Multiplicity | 6.6 (6.7) | | |
| $\langle I/\sigma(I) \rangle$ | 19.5 (11.2) | 8.7 (1.3) | 13.9 (5.5) |
| $CC_{1/2}$ | 0.998 (0.993) | 0.719 (0.547) | 0.997 (0.951) |
| **Refinement statistics** | | | |
| $R_{work}/R_{free}$ (%) | | 21.1/24.4 | 24.3/30.8 |
| Mean protein/ligands/solvent isotropic $B$ factor (Å$^2$) | | 39.0/34.0/41.7 | 62.7/56.2/40.0 |
| Protein assembly in asymmetric unit (AU) | | 4 monomers | 8 monomers |
| Protein residues in gene | | 326 | 328 |
| Total modelled residues in AU | | | |
| - protein residues by chain | | A-B:326,C:325,D:324 | A-F,H:323,G:263 |
| - ligands | | 4 FAD | 8 FAD, 3 NADPH |
| - added waters | | 216 | 26 |
| Matthews coefficient (Å$^3$/Da) | | 2.26 | 2.80 |
| Solvent content (%) | | 45.1 | 55.8 |
| Ramachandran favored/ allowed/outliers (%) | | 96.8/3.0/0.2 | 91.9/6.6/1.5 |
| RMSD bond lengths (Å) | | 0.007 | 0.009 |
| RMSD bond angles (°) | | 0.87 | 1.12 |
| PDB ID | | 7A76 | 7A7B |

**Table S2**: The structures with the highest Z-score for each of the five most similar protein types from the DALI search of the full PDB of *Bc* YpdA.

| Protein | PDBid | RMSD | Z-score | Aligned residues | Number of residues | % sequence identity | Organism |
|---|---|---|---|---|---|---|---|
| FPMO | 4C5O | 2.8 | 28.9 | 289 | 330 | 19 | *Stenotrophomonas maltophilia* |
| TrxR | 5VT3 | 2.4 | 28.1 | 295 | 313 | 19 | *Vibrio vulnificus* |
| DLD | 6CMZ | 3.3 | 25.0 | 285 | 459 | 18 | *Burkholderia cenocepacia* |
| FNR | 3AB1 | 4.6 | 24.9 | 294 | 318 | 22 | *Chlorobaculum tepidum* |
| GR | 1GRB | 3.3 | 23.9 | 278 | 461 | 19 | *Homo sapiens* |

### References

(1) Lofstad, M.; Gudim, I.; Hammerstad, M.; Røhr, Å. K.; Hersleth, H.-P. Activation of the Class Ib Ribonucleotide Reductase by a Flavodoxin Reductase in *Bacillus cereus*. *Biochemistry* **2016,** *55*, 4998-5001.

(2) Sharma, S. V.; Jothivasan, V. K.; Newton, G. L.; Upton, H.; Wakabayashi, J. I.; Kane, M. G.; Roberts, A. A.; Rawat, M.; La Clair, J. J.; Hamilton, C. J. Chemical and Chemoenzymatic Syntheses of Bacillithiol: A Unique Low-Molecular-Weight Thiol Amongst Low G+C Gram-Positive Bacteria. *Angew. Chem. Int. Ed.* **2011,** *50*, 7101-7104.

(3) Mueller, U.; Thunnissen, M.; Nan, J.; Eguiraun, M.; Bolmsten, F.; Milán-Otero, A.; Guijarro, M.; Oscarsson, M.; Dowd, A.; de Sanctis, D.; Leonard, G. MXCuBE3: A New Era of MX-Beamline Control Begins. *Synchrotron Radiat. News* **2017,** *30*, 20-27.

(4) Delageniere, S.; Brenchereau, P.; Launer, L.; Ashton, A. W.; Leal, R.; Veyrier, S.; Gabadinho, J.; Gordon, E. J.; Jones, S. D.; Levik, K. E.; McSweeney, S. M.; Monaco, S.; Nanao, M.; Spruce, D.; Svensson, O.; Walsh, M. A.; Leonard, G. A. ISPyB: An Information Management System for Synchrotron Macromolecular Crystallography. *Bioinformatics* **2011,** *27*, 3186-3192.

(5) Battye, T. G. G.; Kontogiannis, L.; Johnson, O.; Powell, H. R.; Leslie, A. G. W. iMOSFLM: A New Graphical Interface for Diffraction-Image Processing with MOSFLM. *Acta Crystallogr., Sect. D: Biol. Crystallogr.* **2011,** *67*, 271-281.

(6) Incardona, M. F.; Bourenkov, G. P.; Levik, K.; Pieritz, R. A.; Popov, A. N.; Svensson, O. Edna: A Framework for Plugin-Based Applications Applied to X-Ray Experiment Online Data Analysis. *J. Synchrot. Radiat.* **2009,** *16*, 872-879.

(7) Kabsch, W. Xds. *Acta Crystallogr., Sect. D: Biol. Crystallogr.* **2010,** *66*, 125-132.

(8) Vonrhein, C.; Flensburg, C.; Keller, P.; Sharff, A.; Smart, O.; Paciorek, W.; Womack, T.; Bricogne, G. Data Processing and Analysis with the autoPROC Toolbox. *Acta Crystallogr., Sect. D: Biol. Crystallogr.* **2011,** *67*, 293-302.

(9) Winn, M. D.; Ballard, C. C.; Cowtan, K. D.; Dodson, E. J.; Emsley, P.; Evans, P. R.; Keegan, R. M.; Krissinel, E. B.; Leslie, A. G. W.; McCoy, A.; McNicholas, S. J.; Murshudov, G. N.; Pannu, N. S.; Potterton, E. A.; Powell, H. R.; Read, R. J.; Vagin, A.; Wilson, K. S. Overview of the CCP4 Suite and Current Developments. *Acta Crystallogr., Sect. D: Biol. Crystallogr.* **2011,** *67*, 235-242.

(10) Matthews, B. W. Solvent Content of Protein Crystals. *J. Mol. Biol.* **1968,** *33*, 491-&.

(11) Skubak, P.; Pannu, N. S. Automatic Protein Structure Solution from Weak X-Ray Data. *Nat. Commun.* **2013,** *4*.

(12) Sheldrick, G. M. A Short History of Shelx. *Acta Crystallogr., Sect. A* **2008,** *64*, 112-122.

(13) Schneider, T. R.; Sheldrick, G. M. Substructure Solution with SHELXD. *Acta Crystallogr., Sect. D: Biol. Crystallogr.* **2002,** *58*, 1772-1779.

(14) Murshudov, G. N.; Skubak, P.; Lebedev, A. A.; Pannu, N. S.; Steiner, R. A.; Nicholls, R. A.; Winn, M. D.; Long, F.; Vagin, A. A. REFMAC5 for the Refinement of Macromolecular Crystal Structures. *Acta Crystallogr., Sect. D: Biol. Crystallogr.* **2011,** *67*, 355-367.

(15) Abrahams, J. P.; Leslie, A. G. W. Methods Used in the Structure Determination of Bovine Mitochondrial F-1 ATPase. *Acta Crystallogr., Sect. D: Biol. Crystallogr.* **1996,** *52*, 30-42.

(16) Skubak, P.; Waterreus, W. J.; Pannu, N. S. Multivariate Phase Combination Improves Automated Crystallographic Model Building. *Acta Crystallogr., Sect. D: Biol. Crystallogr.* **2010,** *66*, 783-788.
(17) Cowtan, K. Recent Developments in Classical Density Modification. *Acta Crystallogr., Sect. D: Biol. Crystallogr.* **2010,** *66*, 470-478.
(18) Cowtan, K. The Buccaneer Software for Automated Model Building. 1. Tracing Protein Chains. *Acta Crystallogr., Sect. D: Biol. Crystallogr.* **2006,** *62*, 1002-1011.
(19) Langer, G.; Cohen, S. X.; Lamzin, V. S.; Perrakis, A. Automated Macromolecular Model Building for X-Ray Crystallography Using ARP/wARP Version 7. *Nat. Protoc.* **2008,** *3*, 1171-1179.
(20) Perrakis, A.; Morris, R.; Lamzin, V. S. Automated Protein Model Building Combined with Iterative Structure Refinement. *Nat. Struct. Biol.* **1999,** *6*, 458-463.
(21) Perrakis, A.; Harkiolaki, M.; Wilson, K. S.; Lamzin, V. S. ARP/wARP and Molecular Replacement. *Acta Crystallogr., Sect. D: Biol. Crystallogr.* **2001,** *57*, 1445-1450.
(22) Afonine, P. V.; Grosse-Kunstleve, R. W.; Echols, N.; Headd, J. J.; Moriarty, N. W.; Mustyakimov, M.; Terwilliger, T. C.; Urzhumtsev, A.; Zwart, P. H.; Adams, P. D. Towards Automated Crystallographic Structure Refinement with Phenix.Refine. *Acta Crystallogr., Sect. D: Biol. Crystallogr.* **2012,** *68*, 352-367.
(23) Adams, P. D.; Afonine, P. V.; Bunkoczi, G.; Chen, V. B.; Davis, I. W.; Echols, N.; Headd, J. J.; Hung, L. W.; Kapral, G. J.; Grosse-Kunstleve, R. W.; McCoy, A. J.; Moriarty, N. W.; Oeffner, R.; Read, R. J.; Richardson, D. C.; Richardson, J. S.; Terwilliger, T. C.; Zwart, P. H. Phenix: A Comprehensive Python-Based System for Macromolecular Structure Solution. *Acta Crystallogr., Sect. D: Biol. Crystallogr.* **2010,** *66*, 213-221.
(24) Emsley, P.; Lohkamp, B.; Scott, W. G.; Cowtan, K. Features and Development of Coot. *Acta Crystallogr., Sect. D: Biol. Crystallogr.* **2010,** *66*, 486-501.
(25) Chen, V. B.; Arendall, W. B.; Headd, J. J.; Keedy, D. A.; Immormino, R. M.; Kapral, G. J.; Murray, L. W.; Richardson, J. S.; Richardson, D. C. MolProbity: All-Atom Structure Validation for Macromolecular Crystallography. *Acta Crystallogr., Sect. D: Biol. Crystallogr.* **2010,** *66*, 12-21.
(26) McCoy, A. J.; Grosse-Kunstleve, R. W.; Adams, P. D.; Winn, M. D.; Storoni, L. C.; Read, R. J. Phaser Crystallographic Software. *J. Appl. Crystallogr.* **2007,** *40*, 658-674.
(27) Røhr, Å. K.; Hersleth, H.-P.; Andersson, K. K. Tracking Flavin Conformations in Protein Crystal Structures with Raman Spectroscopy and QM/MM Calculations. *Angew. Chem. Int. Ed.* **2010,** *49*, 2324-2327.
(28) Hiras, J.; Sharma, S. V.; Raman, V.; Tinson, R. A. J.; Arbach, M.; Rodrigues, D. F.; Norambuena, J.; Hamilton, C. J.; Hanson, T. E. Physiological Studies of Chlorobiaceae Suggest That Bacillithiol Derivatives Are the Most Widespread Thiols in Bacteria. *mBio* **2018,** *9*.
(29) Waterhouse, A. M.; Procter, J. B.; Martin, D. M. A.; Clamp, M.; Barton, G. J. Jalview Version 2-a Multiple Sequence Alignment Editor and Analysis Workbench. *Bioinformatics* **2009,** *25*, 1189-1191.
(30) Sievers, F.; Wilm, A.; Dineen, D.; Gibson, T. J.; Karplus, K.; Li, W. Z.; Lopez, R.; McWilliam, H.; Remmert, M.; Soding, J.; Thompson, J. D.; Higgins, D. G. Fast, Scalable Generation of High-Quality Protein Multiple Sequence Alignments Using Clustal Omega. *Mol. Syst. Biol.* **2011,** *7*.
(31) Linzner, N.; Loi, V. V.; Fritsch, V. N.; Tung, Q. N.; Stenzel, S.; Wirtz, M.; Hell, R.; Hamilton, C. J.; Tedin, K.; Fulde, M.; Antelmann, H. *Staphylococcus aureus* Uses the Bacilliredoxin (BrxAB)/Bacillithiol Disulfide Reductase (YpdA) Redox Pathway to Defend against Oxidative Stress under Infections. *Front. Microbiol.* **2019,** *10*.

(32) Holm, L. Benchmarking Fold Detection by DaliLite V.5. *Bioinformatics* **2019,** *35*, 5326-5327.
(33) Touw, W. G.; Baakman, C.; Black, J.; te Beek, T. A. H.; Krieger, E.; Joosten, R. P.; Vriend, G. A Series of PDB-Related Databanks for Everyday Needs. *Nucleic Acids Res.* **2015,** *43*, D364-D368.
(34) Kabsch, W.; Sander, C. Dictionary of Protein Secondary Structure - Pattern-Recognition of Hydrogen-Bonded and Geometrical Features. *Biopolymers* **1983,** *22*, 2577-2637.
(35) Landau, M.; Mayrose, I.; Rosenberg, Y.; Glaser, F.; Martz, E.; Pupko, T.; Ben-Tal, N. ConSurf 2005: The Projection of Evolutionary Conservation Scores of Residues on Protein Structures. *Nucleic Acids Res.* **2005,** *33*, W299-W302.
(36) Ashkenazy, H.; Erez, E.; Martz, E.; Pupko, T.; Ben-Tal, N. ConSurf 2010: Calculating Evolutionary Conservation in Sequence and Structure of Proteins and Nucleic Acids. *Nucleic Acids Res.* **2010,** *38*, W529-W533.
(37) Glaser, F.; Pupko, T.; Paz, I.; Bell, R. E.; Bechor-Shental, D.; Martz, E.; Ben-Tal, N. Consurf: Identification of Functional Regions in Proteins by Surface-Mapping of Phylogenetic Information. *Bioinformatics* **2003,** *19*, 163-164.
(38) Celniker, G.; Nimrod, G.; Ashkenazy, H.; Glaser, F.; Martz, E.; Mayrose, I.; Pupko, T.; Ben-Tal, N. Consurf: Using Evolutionary Data to Raise Testable Hypotheses About Protein Function. *Isr. J. Chem.* **2013,** *53*, 199-206.
(39) Ho, B. K.; Gruswitz, F. HOLLOW: Generating Accurate Representations of Channel and Interior Surfaces in Molecular Structures. *BMC Struct. Biol.* **2008,** *8*, 49.
(40) Wallace, A. C.; Laskowski, R. A.; Thornton, J. M. Ligplot - a Program to Generate Schematic Diagrams of Protein Ligand Interactions. *Protein Eng.* **1995,** *8*, 127-134.
(41) Laskowski, R. A.; Swindells, M. B. LigPlot$^{+}$: Multiple Ligand-Protein Interaction Diagrams for Drug Discovery. *J. Chem. Inf. Model* **2011,** *51*, 2778-2786.
